# Insight into the regulatory mechanism of the MFS transporter, SCO4121 by the MarR regulator, SCO4122

**DOI:** 10.1101/2024.02.28.577416

**Authors:** Ankita Nag, Shiksha Sharma, Pittu Sandhya Rani, Sarika Mehra

**Author notes:** Ankita Nag: Department of Microbiology and Immunology, Emory University School of Medicine, Atlanta, Georgia.

## Abstract

MarR group of transcriptional regulators are ubiquitous in bacteria and found to be involved in regulation of efflux pumps that confer multidrug resistance phenotype. While most characterized MarR regulators act as transcriptional repressors, we earlier identified a MarR regulator SCO4122 in *Streptomyces coelicolor*, playing an essential role in transcriptional activation of the MFS transporter SCO4121 in response to multiple substrates of the latter, including streptomycin, ciprofloxacin and chloramphenicol. In this study, using Surface Plasmon Resonance, we demonstrate that SCO4122 interacts directly with the diverse substrates of SCO4121 with the highest affinity for streptomycin with a KD of 0.73 μM. Further, in-vitro and in-vivo studies reveal that SCO4122 also binds to the intergenic region between *sco4121* and *sco4122,* where the interaction is dependent on the cooperative binding of SCO4122 to three motifs in this region. A conserved Methionine, M93, in SCO4122 is identified to be an integral amino acid residue that is involved in activation of SCO4121 in response to ciprofloxacin, streptomycin and EtBr but not chloramphenicol. Furthermore, our studies also indicate that upon binding to different substrates, the affinity of SCO4122 to the *sco4121* promoter increases 50-1000 fold, thereby leading to enhanced expression of the transporter, SCO4121. This study thus highlights that SCO4122 is a novel MarR regulator that functions as a strong transcriptional activator of an efflux pump, SCO4121, through intricate molecular mechanisms in presence of structurally dissimilar substrates.

**Graphical abstract:** 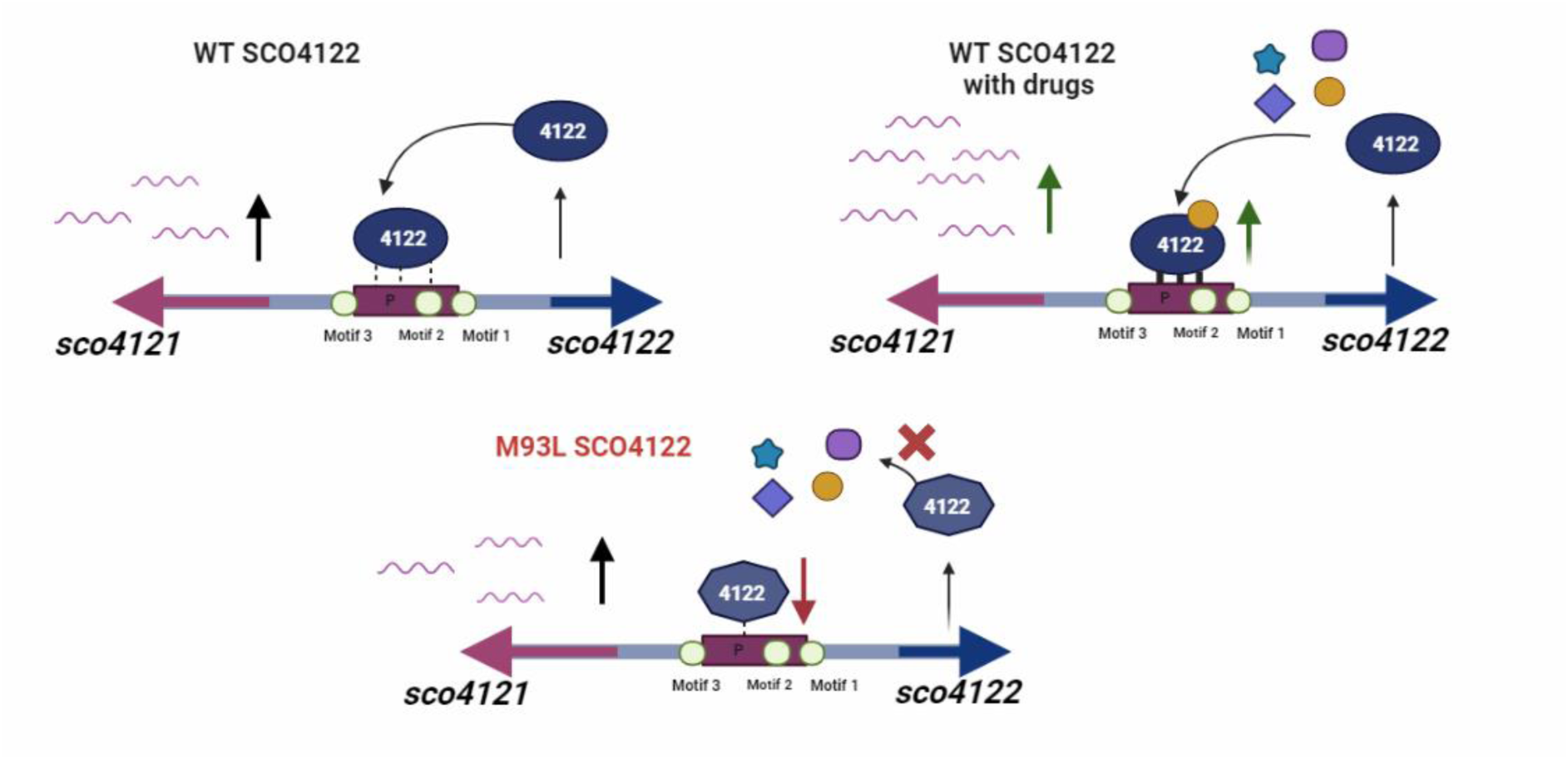

## Introduction

Expression of multidrug efflux pumps is tightly regulated by either transcriptional repressors or activators. In majority of the cases, the regulators are induced in response to the chemically diverse compounds that the efflux pumps recognize as substrates. These regulators are cytosolic in nature and thus act as sensors, responding to the toxic compounds that the bacteria sense under different environmental conditions. Both two-components and one-component regulators are involved in regulation of several efflux pumps in bacteria (1, 2). For example, the two-component system RocS2-RocA2 acts as a repressor of the MexAB-OprM efflux pump whereas, ParR-ParS upregulates the expression of MexEF-OprN through MexS in *Pseudomonas aeruginosa* (3, 4). Similarly, the two-component system SmeRySy is involved in negative regulation of the efflux pump SmeDEF, which provides multidrug resistance in the pathogen *Stenotrophomonas maltophilia* (5). The one-component regulatory systems form a dominant part of various signaling events in bacteria and majority of the efflux pumps, are found to be regulated by these systems (6). These regulators are composed of a DNA-binding domain consisting of a Helix-Turn-Helix (HTH) motif and a sensory domain which is involved in protein oligomerization. The activity of these regulators is generally controlled by binding to diverse compounds (7). The MarR, TetR, LacI, MerR and LysR family of proteins are the major one-component systems ubiquitously distributed in different bacteria (8–12).

MarR (Multiple Antibiotic Resistance) regulators are ubiquitously distributed in different Gram-positive and Gram-negative bacteria that operate to differentially regulate several efflux systems. Majority of the MarR regulators are transcriptional repressors of the efflux genes. One such widely studied MarR regulator is the *E. coli* MarR protein which represses the expression of the multidrug resistant efflux pump AcrAB (13, 14). The FarR in *Neisseria gonorrhoeae*, MexR in *Pseudomonas aeruginosa*, and the MepR *Staphylococcus aureus* are some of the other widely studied MarR regulators that regulate the multidrug efflux systems MexAB-OprM, FarAB and MepA respectively via transcriptional repression (15–17). However, very few MarR regulators have been found that are involved in transcriptional activation of transporter genes. One such example is BldR which regulates the multidrug transporter Sso1351 in the archaea *Sulfolobus solfatiricus* (18). The PenR in *Streptomyces exfolitus* is another example of a MarR regulator that positively regulates the genes involved in the export of the sesquiterpenoid antibiotic pentalenolactone (19).

MarR regulators have been reported to interact with a wide array of compounds, thereby allowing them to function under variable environments. For example, phenolic compounds such as salicylate were shown to cause envelope stress, resulting in release of copper ions, which are actual ligands for E.coli MarR (20). Similarly, the MarR regulator MprA (also previously known as EmrR) in *E. coli* interacts with compounds such as carbonyl cyanide *m*-chlorophenyldrazone, 2,4-dinitrophenol, and nalidixic acid, thereby allowing derepression and subsequent activation of the efflux pump EmrAB (21). A recent study has also identified a MarR repressor, MhqR in *Staphylococcus aureus* that provides resistance to fluoroquinolone drugs via direct binding of the quinolone molecule (22). MarR regulators have also been found to bind to more than one site in the promoter of the cognate genes that they regulate. For example, the FarR regulator in *N. gonorrhoeae* binds to two sites upstream of FarAB efflux pump to modulate its expression (23). Similarly, a cooperative binding to two sites between −35 and −10 promoter sequences by MepR results in repression of the MepA efflux pump in *Staphylococcus aureus* (24, 25).

Previous studies in our lab have identified a MarR regulator, SCO4122 to be associated in positive regulation of the MFS transporter, SCO4121 in presence of different substrates of SCO4121. In-vivo studies revealed this regulation to be direct without involvement of any additional regulatory proteins (26). In this study, we have evaluated the detailed molecular mechanism by which SCO4122 regulates the expression of SCO4121 in response to different drugs. We found that the substrates of SCO4121 interact directly with SCO4122, thereby resulting in increased affinity of SCO4122 to the *sco4121* promoter and increased expression of SCO4121. Additionally, we have also highlighted the importance of a conserved methionine residue M93 in SCO4122 to play an essential role in activation of SCO4122 with subsequent activation of SCO4121.

## Results

### Activation of SCO4121 occurs via binding of SCO4122 to specific sequences in the *sco4121* promoter

SCO4122 encodes a 26 KDa protein belonging to the MarR group of transcriptional regulators. Previously through in-vivo studies, we have shown SCO4122 to be involved in direct activation of the efflux pump SCO4121 in presence of different drugs that are recognized as substrates of the latter (26). In this study, we investigate the biophysical mechanism of this activation through an in-vitro approach using Surface Plasmon Resonance (SPR). SCO4122 is hypothesized to activate the expression of SCO4121 by binding to the DNA region between *sco4121* and *sco4122* (*sco4121* control region). We measured the affinity of interaction between SCO4122 and the *sco4121* control region consisting of the putative *sco4121* promoter. SCO4122 protein was purified to homogeneity and immobilized to the CM5 sensor chip through amine coupling. The DNA region consisting the 5’ to 3’ fragment of the *sco4121* promoter was passed over the purified SCO4122 protein at different concentrations and affinity was calculated by measuring the K_D_ of interaction. We found that SCO4122 was able to directly bind to the DNA segment bearing the *sco4121* promoter with an equilibrium dissociation constant (K_D_) of 0.4 µM (Figure 1A). Negligible binding was observed when a non-specific DNA sequence was passed over the immobilized SCO4122 protein (Figure S1), thereby suggesting sequence specific binding of SCO4122 to the *sco4121* promoter. The details of the sequence of the *sco4121* promoter region and the non-specific DNA are presented in Table S1.

**Figure 1:**
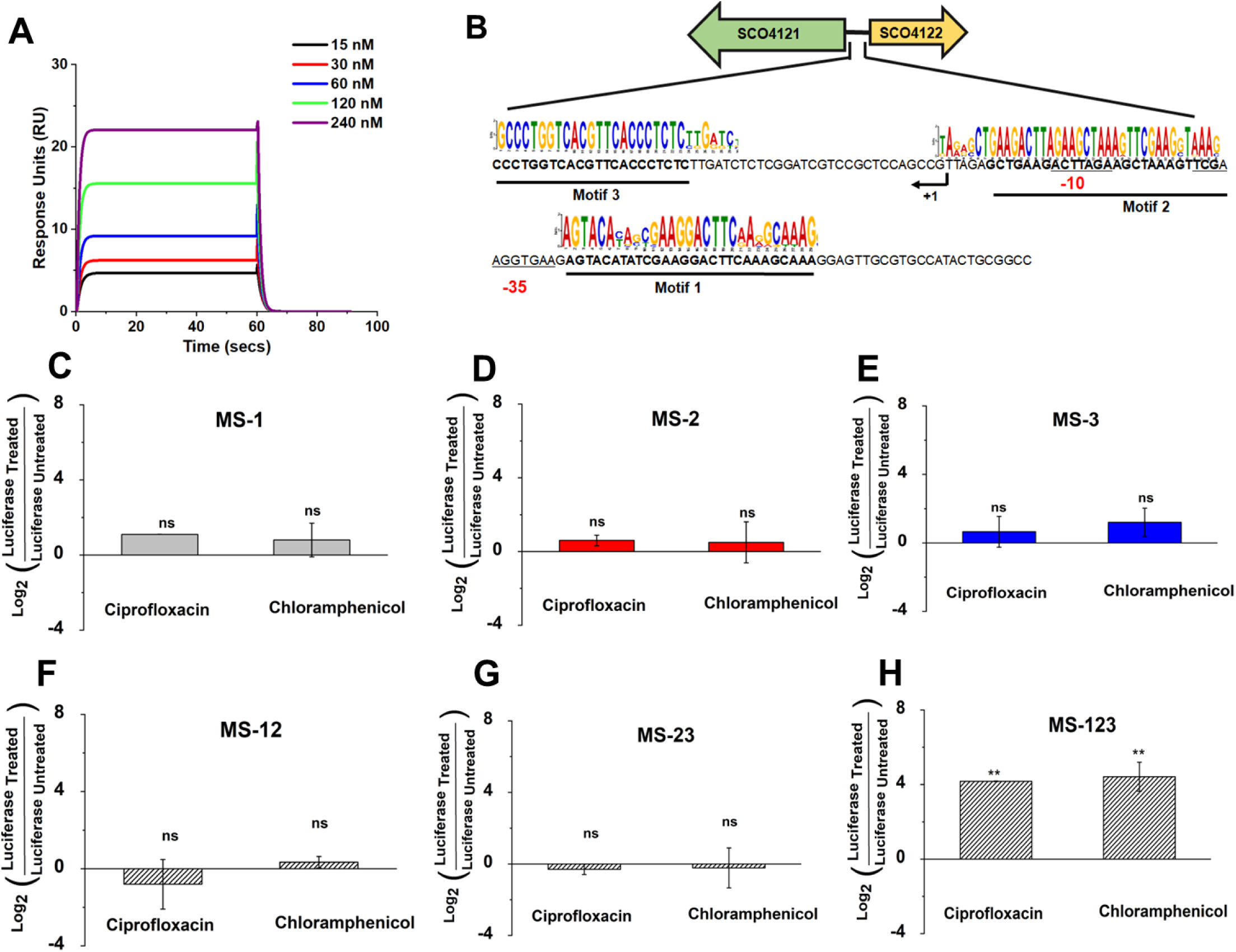
Estimation of binding of SCO4122 to the *sco4121* promoter using in-vitro and in-vivo studies. (A) Concentration dependent binding of *sco4121* promoter to the purified SCO4122 protein measured using SPR assay. *sco4121* promoter bound to the SCO4122 protein in a concentration dependent manner with a K_D_ of 0.4 µM. (B) Schematic showing the intergenic region between *sco4121* and s*co4122* and placement of the three motifs (highlighted in bold) predicted to be binding sites of SCO4122. Underlined bases depict −10 and −35 of the promoter region of *sco4121*. The motifs were discovered using the MEME tool by aligning of 30 Streptomyces species, where homologs of SCO4121 and SCO4122 were adjacently present with an identity of greater than 90%. Promoter regions were predicted using BPROM tool. (C) Expression of the luciferase gene measured under inhibitory concentrations of ciprofloxacin (0.4 µg/ml) and chloramphenicol (7 µg/ml) in MS-1 cells, bearing only Motif 1. (D) Expression of the luciferase gene measured under inhibitory concentrations of ciprofloxacin (0.4 µg/ml) and chloramphenicol (7 µg/ml) in MS-1 cells, bearing only Motif 2 (E) Expression of the luciferase gene measured under inhibitory concentrations of ciprofloxacin (0.4 µg/ml) and chloramphenicol (7 µg/ml) in MS-1 cells, bearing only Motif 3 (F) Expression of the luciferase gene measured under inhibitory concentrations of ciprofloxacin (0.4 µg/ml) and chloramphenicol (7 µg/ml) in MS-1 cells, bearing Motif 1 and Motif 2. (G) Expression of the luciferase gene measured under inhibitory concentrations of ciprofloxacin (0.4 µg/ml) and chloramphenicol (7 µg/ml) in MS-1 cells, bearing Motif 2 and Motif 3. (H) Expression of the luciferase gene measured under inhibitory concentrations of ciprofloxacin (0.4 µg/ml) and chloramphenicol (7 µg/ml) in MS-1 cells, bearing Motif 1, Motif 2 and Motif 3 together. Expression of luciferase could be observed in only MS123 cells, where all the motifs (Motif 1, Motif 2 and Motif 3) were intact. Expression was measured using Real-Time qRT PCR and normalized with 16S as the house-keeping gene followed by normalization with untreated cells. Error bars calculated with respect to three biological triplicates. ** indicates p-value < 0.005, ns indicates non-significant.

To further unravel the region where SCO4122 binds in the intergenic region between *sco4121* and *sco4122* (*sco4121* control region), we utilized the MEME program. Using this tool, we identified three motifs in this region, which were predicted to be the plausible sites of binding of SCO4122. These motifs were discovered by aligning intergenic regions of *Streptomyces* species where homologs of both SCO4121 and SCO4122 were present adjacently and conserved with an identity of >90% (26). The three motifs were found to be about 26±2 bp in size and non-palindromic in nature (Figure 1B). The distribution and conservation of these three motifs in some Streptomyces has been depicted in Figure S2. To determine which of these specific motifs are essential for binding of SCO4122, we measured the promoter activity of *sco4121* by preparing different constructs with the three motifs under a downstream luciferase gene, shown in Figure S3. These constructs were individually incorporated into *M. smegmatis* MS4122 cells, which constitutively expressed the *sco4122* gene. Expression of luciferase mRNA was monitored in these cells in presence of ciprofloxacin and chloramphenicol, the two major substrates of SCO4121 (26).When subjected to inhibitory concentrations of the drugs, increased luciferase expression was observed in those cells where all the three motifs (Motif 1, 2 and 3) were intact (Figure 1C). However, no luciferase expression was observed when the motifs were present either individually (Figure 1D, 1E and 1F) or in a combination of two (Figure 1G and 1H). MarR regulators are dimeric in nature and mostly bind to short palindromic sequences in the promoter of the target gene (27). In many cases, cooperative binding has been seen where one dimer binds to one motif, whereas the second dimer binds to the other motif such as the MarR regulator OhrR in *Bacillus subtillis* (28). Similar phenomena has been noticed in case of the MepR regulator in *Staphylococcus aureus,* where binding of subsequent half sites results in cooperative binding, thus leading to repression of the target gene (29). On similar lines, we found that SCO4121 activity is dependent on the presence of all the three motifs, and an involvement of a possible cooperative binding of SCO4122 to these sites results in increased expression of *sco4121* in presence of the drugs.

### SCO4122 directly interacts with multiple drugs to activate expression of SCO4121

Our previous study suggests that SCO4121 is induced in response to its diverse substrates in presence of SCO4122 (26). In order to investigate whether the substrates of SCO4121 directly interact with SCO4122 to activate the expression of the former, an in-vitro binding study of the drugs to the SCO4122 protein was performed. This study was done using the SPR assay, where drugs at varying concentrations were passed over the already immobilized SCO4122 protein to the CM5 sensor chip. The interaction was further monitored by calculating the equilibrium dissociation constant (K_D_). We find that SCO4122 was able to bind to the substrates of SCO4121 in a concentration dependent manner (Figure 2). Streptomycin bound to the SCO4122 protein with the highest affinity, followed by EtBr and ciprofloxacin with a K_D_ of 0.7 µM, 1. 39 µM and 31 µM respectively. Interestingly, chloramphenicol, a major substrate of SCO4121 showed a weak binding to SCO4122 with a K_D_ of 2000 µM (Table 1). Curve-fitting data further revealed that all drugs bound to the SCO4122 protein as per the 1:1 Langmuir stoichiometry model. However, chloramphenicol on the other hand fit into the Two-state binding model, thus suggesting a 2-step binding of the drug to the SCO4122 protein. It was interesting that tetracycline, which was not recognized as a substrate of SCO4121, also showed binding to the SCO4122 protein, with K_D_ value of 2.68 µM (Table 1). Thus, SCO4122 can bind to a cohort of drugs with varying affinities, suggesting that this MarR regulator can function in response to variable, and rapidly changing environments.

**Figure 2:**
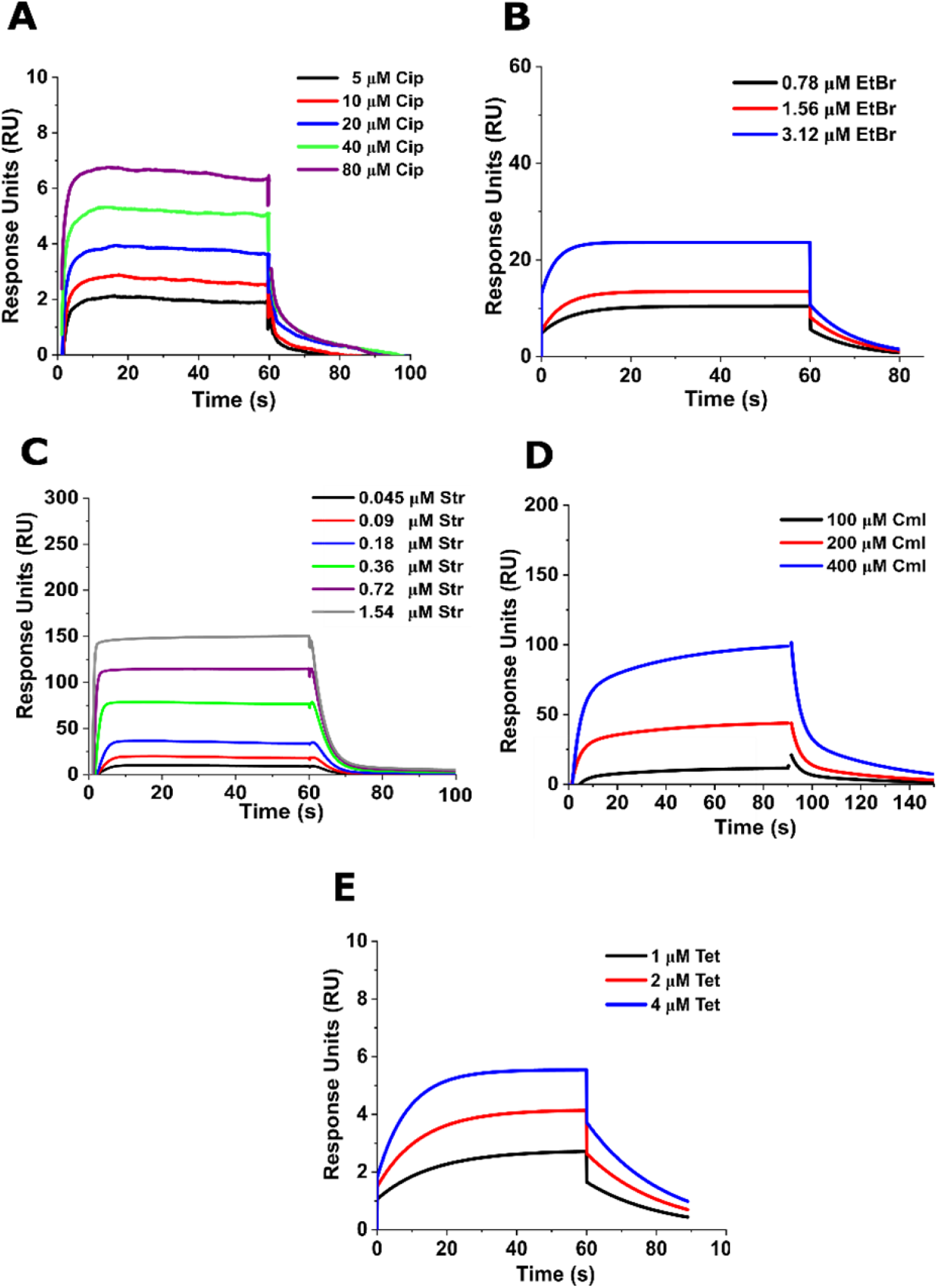
Binding kinetics of different drugs to SCO4122 protein measured via SPR assay (A) Ciprofloxacin (B) EtBr (C) Streptomycin (D) Chloramphenicol (E) Tetracycline. SCO4122 was immobilized over CM5 Sensor chip through Amide linkage. All drugs (ciprofloxacin, EtBr, streptomycin, chloramphenicol and tetracycline) were passed over the immobilized SCO4122 protein at a concentration dependent manner. The Equilibrium Dissociation constant (K_D_) is calculated using the Biacore software. Ciprofloxacin bound to SCO4122 with K_D_ of 31 µM, EtBr bound to SCO4122 with a K_D_ of 1.39 µM, streptomycin bound to SCO4122 with a K_D_ of 0.73 µM, chloramphenicol bound to SCO4122 with a K_D_ of 2000 µM and tetracycline bound to SCO4122 with a K_D_ of 2.68 µM. Binding of all drugs with SCO4122 was studied using an association time of 60 seconds followed by a dissociation time of 120 seconds.

**Table 1:** Equilibrium Dissociation constant (K_D_) of DNA and different drugs to the WT and M93L mutant SCO4122 protein calculated using Surface Plasmon Resonance method.

| Drugs | Equilibrium Dissociation constant ($K_D$) (μM) | |
| --- | --- | --- |
|  | WT | M93L |
| Only DNA | 0.4 | 2.11 |
| Ciprofloxacin | 31 | NA |
| EtBr | 1.39 | NA |
| Streptomycin | 0.73 | 53.9 |
| Chloramphenicol | 2000 | NA |
| Tetracycline | 2.68 | 2.43 |

Note: $K_D$ for ciprofloxacin, EtBr, Streptomycin and EtBr calculated by fitting curve as per 1:1 Langmuir stoichiometry. $K_D$ for chloramphenicol was calculated by fitting curve into the Two-State binding stoichiometry.

### A conserved Methionine M93 is essential for activation of SCO4121 by SCO4122

As direct interaction of drugs with SCO4122 was noted from our previous results, we further aimed to decipher the amino acid residue in the SCO4122 protein that potentiates the activation of SCO4121 in response to the diverse drugs. Using BLASTp analysis, we identified SCO4122 to be ubiquitously distributed among *Streptomyces* isolated from different environmental niche (Figure S4). Apart from homologs in *Streptomyces*, SCO4122 was found to be distributed in other soil bacteria. Among the different conserved amino acids, we found a conserved methionine M93 in all the homologs (Figure S4). Previous studies have identified the role of methionine in active site of many proteins, both in prokaryotes and eukaryotes, to be essential in their activity. In the efflux pump CusA, encoded by the *E. coli* genome, methionine aids in binding to different metals (30). Methionine has also been found to be a conserved residue in the active site of the enzyme beta galactosidase (31). Additionally, methionine has been found to be an integral residue that maintains structural integrity of many proteins. For example, methionine 145 has been found to be critical for dimerization of bovine beta-lactoglobulin (32). This further led us to evaluate whether the conserved Methionine M93 has a role in activation of SCO4122. To test this, we performed a site directed mutagenesis, where the Methionine M93 was replaced by a Leucine residue (Figure 3A), using the *sco4122* gene cloned under the pIJ86 plasmids as the template to amplify the mutated construct. We noted that conversion of the Methionine M93 to Leucine had a least impact on the expression of *sco4122* gene and expression levels of *sco4122* were similar to cΔ4122 strains, which carried the native copies of *sco4122* gene placed under the pIJ86 plasmids, thereby suggesting the conversion of Methionine to Leucine to be a safe substitution (Figure S5). These constructs bearing the M93L mutation were transformed into Δ4122 strains, thus yielding the strain M93L-4122. Using Real Time PCR assay, we further studied the expression of *sco4121* in *S. coelicolor* cells expressing the mutant M93L-SCO4122 protein in presence of different drugs identified as substrates of SCO4121. In presence of MIC and 2X MIC concentrations of ciprofloxacin, EtBr, streptomycin and chloramphenicol, increased expression of *sco4121* was noticed in cΔ4122 strains (Fig 3B-3E). In M93L-4122 strains, basal expression of *sco4121* was found to be similar to that of cΔ4122 cells. However, no further upregulation of *sco4121* was noted upon addition of the drugs ciprofloxacin, EtBr and streptomycin (Figure 3B, 3C, 3D). Contrastingly, *sco4121* levels were found to be upregulated and similar to that of cΔ4122 cells in presence of chloramphenicol in M93L-4122 strains (Figure 3E). This data suggests that the conserved methionine M93 residue probably has a specific role in activation of SCO4122 in response to ciprofloxacin, streptomycin and EtBr. However, an alternate unknown mechanism may exist for activation of *sco4121* in response to chloramphenicol, which may be independent of M93 in SCO4122.

**Figure 3:**
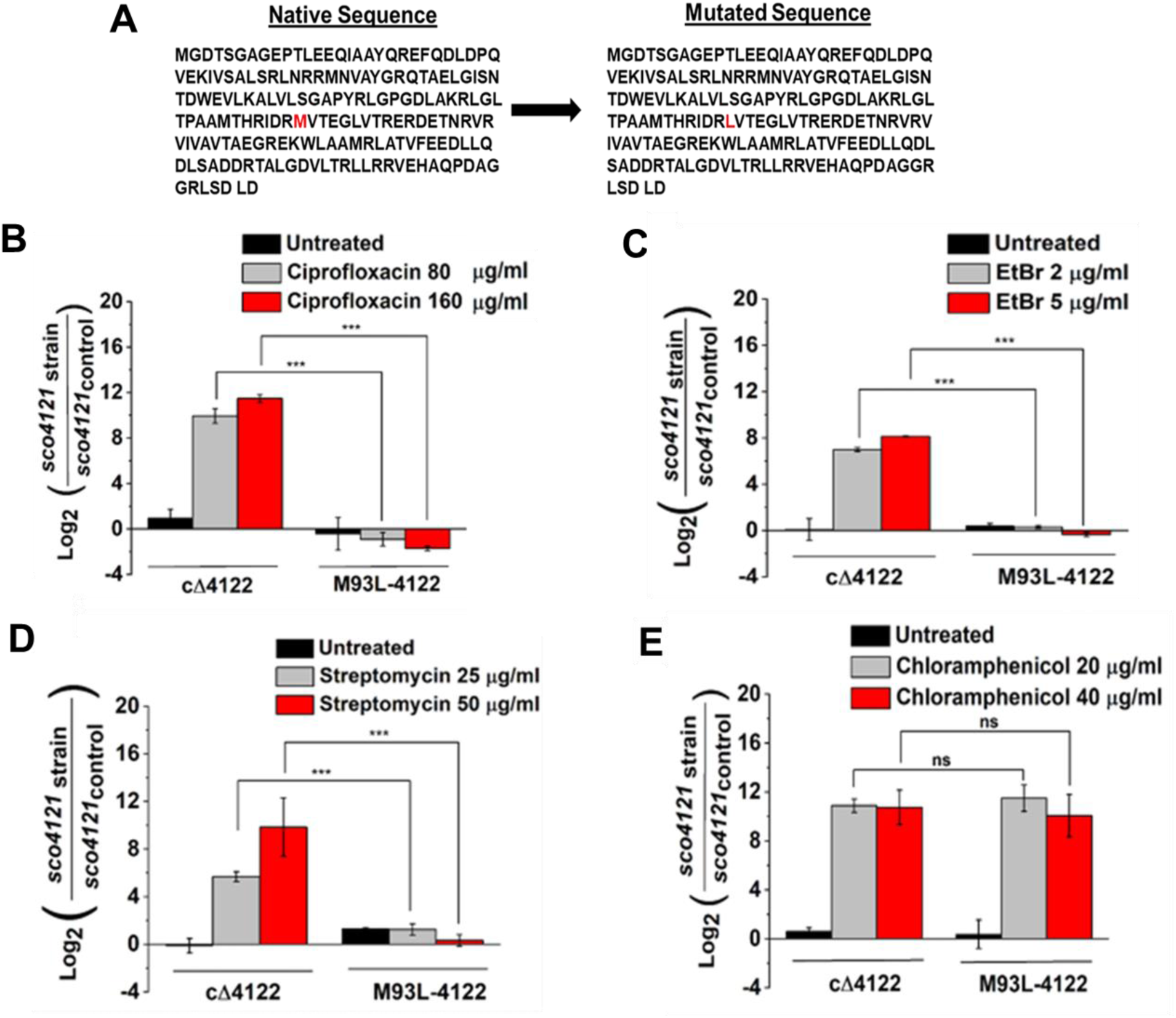
Effect of mutation of Methionine M93 on expression of *sco4121*. (A) Sequence of native and mutated SCO4122 protein. Mutated SCO4122 protein was generated by substitution of Methionine M93 with a Leucine residue. (B) Expression of *sco4121* in cΔ4122 and M93L-4122 cells measured in presence of MIC (80 µg/ml) and 2X MIC (160 µg/ml) concentration of ciprofloxacin (B) Expression of *sco4121* in cΔ4122 and M93L-4122 cells measured in presence of MIC (25 µg/ml) and 2X MIC (50 µg/ml) concentration of streptomycin. (C) Expression of *sco4121* in cΔ4122 and M93L-4122 cells measured in presence of MIC (2 µg/ml) and 2X MIC (5 µg/ml) concentration of EtBr (D) Expression of *sco4121* in cΔ4122 and M93L-4122 cells measured in presence of MIC (20 µg/ml) and 2X MIC (40 µg/ml) concentration of chloramphenicol. *sco4121* levels in M93L-4122 cells were negligible as compared to that observed in cΔ4122 cells in response to ciprofloxacin, EtBr and streptomycin. In presence of chloramphenicol *sco4121* expression was found to be similar between M93L-4122 and cΔ4122 cells. Gene expression was measured using Real Time Quantitative PCR and normalized with 23S as the house keeping gene, followed by normalization with the untreated WT86 (cells harboring empty pIJ86) cells. Statistical significance was calculated with respect to cΔ4122 cells. ***represents p<0.005, ** represents p<0.005, * represents p<0.05, ns represents non-significant

### Differential resistance profile of M93L-4122 strains towards diverse antibiotics

Our studies so far have suggested that many drugs directly interact with SCO4122 leading to activation of SCO4121. We also deciphered the role of a conserved methionine M93 in SCO4122 to be essential in activation of SCO4121. Next, we performed experiments to understand the susceptibility of the M93L-4122 strains towards these different substrates. Previously, we have shown that when Δ4122 cells were complemented with native *sco4122* copies, it led to increased resistance towards different SCO4121 substrates (26). Interestingly, we found that M93L-4122 cells displayed increased susceptibility towards ciprofloxacin, EtBr and streptomycin as compared to the cΔ4122 cells, when tested at different concentrations (Figure 4A, 4B, 4C), thus correlating with the decreased expression of *sco4121* in these cells. This data thus indicates that methionine M93 is a potential site of SCO4122 activation, which further upregulates SCO4121 to provide resistance to these drugs. Interestingly, susceptibility towards chloramphenicol in M93L-4122 cells was similar to that of cΔ4122 cells at higher drug concentrations (Figure 4D). This indicates the involvement of either additional residues beyond M93 or other molecular mechanisms that play an essential part in chloramphenicol resistance in these cells.

**Figure 4:**
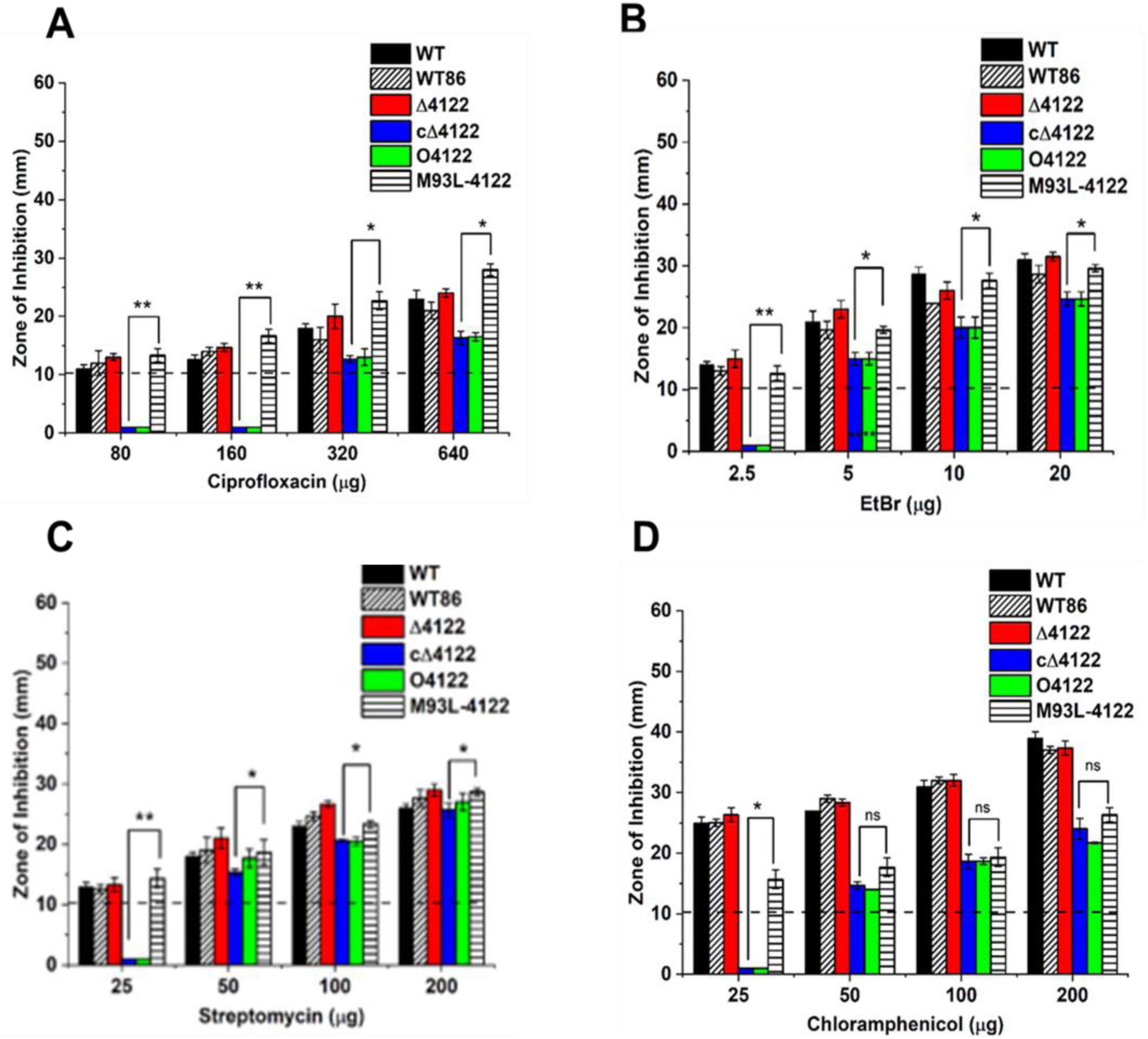
Susceptibility of WT and *S. coelicolor* recombinants WT86, O4122, Δ4122, cΔ4122 and M93L-4122 towards different substrates of SCO4121 (A) ciprofloxacin (B) EtBr (C) streptomycin (D) chloramphenicol. M93L-4122 cells displayed increased susceptibility towards ciprofloxacin, streptomycin and EtBr as compared to cΔ4122 cells. Both M93L-4122 and cΔ4122 cells displayed similar susceptibility profile towards chloramphenicol. Disc Diffusion Assay was used to measure the susceptibility towards the different drugs. Dotted lines represent the minimum zone diameter that can be measured using Disc Diffusion Assay. Statistical significance was calculated with respect to cΔ4122 cells. *** represents p<0.005, ** represents p<0.005, * represents p<0.05, ns represents non-significant.

### Mutation of M93 in SCO4122 eliminates binding to different drugs

Our results so far highlight that ciprofloxacin, streptomycin and EtBr do not lead to increase in expression levels of *sco4121* when the conserved methionine M93 in SCO4122 is mutated. The expression levels of *sco4121* were similar as observed in the WT86 *S. coelicolor* cells in presence of these three drugs (Figure 3). We also observed susceptibility of M93L-4122 strains to different drugs to be comparable to that of the WT86 cells (Figure 4). However, the M93L mutation in SCO4122 did not affect the expression of *sco4121* and the susceptibility profile of M93L-4122 strain in response to chloramphenicol was similar to cΔ4122 cells (Figure 3 and 4). To further understand if the conserved M93 in SCO4122 has a role in direct binding of the different substrates of SCO4121, we measured the interaction of the mutant SCO4122 protein with different drugs using the SPR assay. The mutant M93L SCO4122 protein was purified to homogeneity and immobilized on CM5 sensor chip. The different drugs were individually passed over the mutant protein at varying concentrations and the binding affinity between them was measured by monitoring the K_D_ of interaction. We observed that mutant M93L SCO4122 protein was unable to bind to ciprofloxacin, EtBr and chloramphenicol (Figure 5A, 5B, 5C). Binding of the mutant protein to streptomycin was observed, however with a fairly high K_D_ of 194 µM, which corresponded to nearly 74-folds higher than the K_D_ observed when bound to the native SCO4122 protein (Figure 5D). This data thus suggests that Methionine M93 in SCO4122 forms an essential part of binding to different substrates of SCO4121 and its subsequent activation in response to these drugs. It is interesting to note that the mutant M93L SCO4122 protein was unable to bind to chloramphenicol, despite showing elevated expression of *sco4121* in its presence (Figure 4E). Thus, we observe that under in-vitro conditions methionine M93 is involved in binding of SCO4122 to chloramphenicol, however, activation of SCO4121 by SCO4122 in presence of chloramphenicol may be through an alternate mechanism under in-vivo environments, which could be independent of binding of the drug to the SCO4122 protein. Tetracycline was used a negative control for this assay as it did not induce the expression of *sco4121* (Data not shown). The mutant M93L SCO4122 protein bound to tetracycline with a K_D_ of 2.4 µM, which was similar to the WT SCO4122 protein. The equilibrium dissociation constant of the drugs to the M93L SCO4122 mutant protein is presented in Table 1. Note that the mutant M93L SCO4122 protein bound to *sco4121* control region with a K_D_ of 2.11 µM, which is nearly 5-fold higher than the K_D_ for the WT SCO4122 protein bound to the *sco4121* control region (Figure S6).

**Figure 5:**
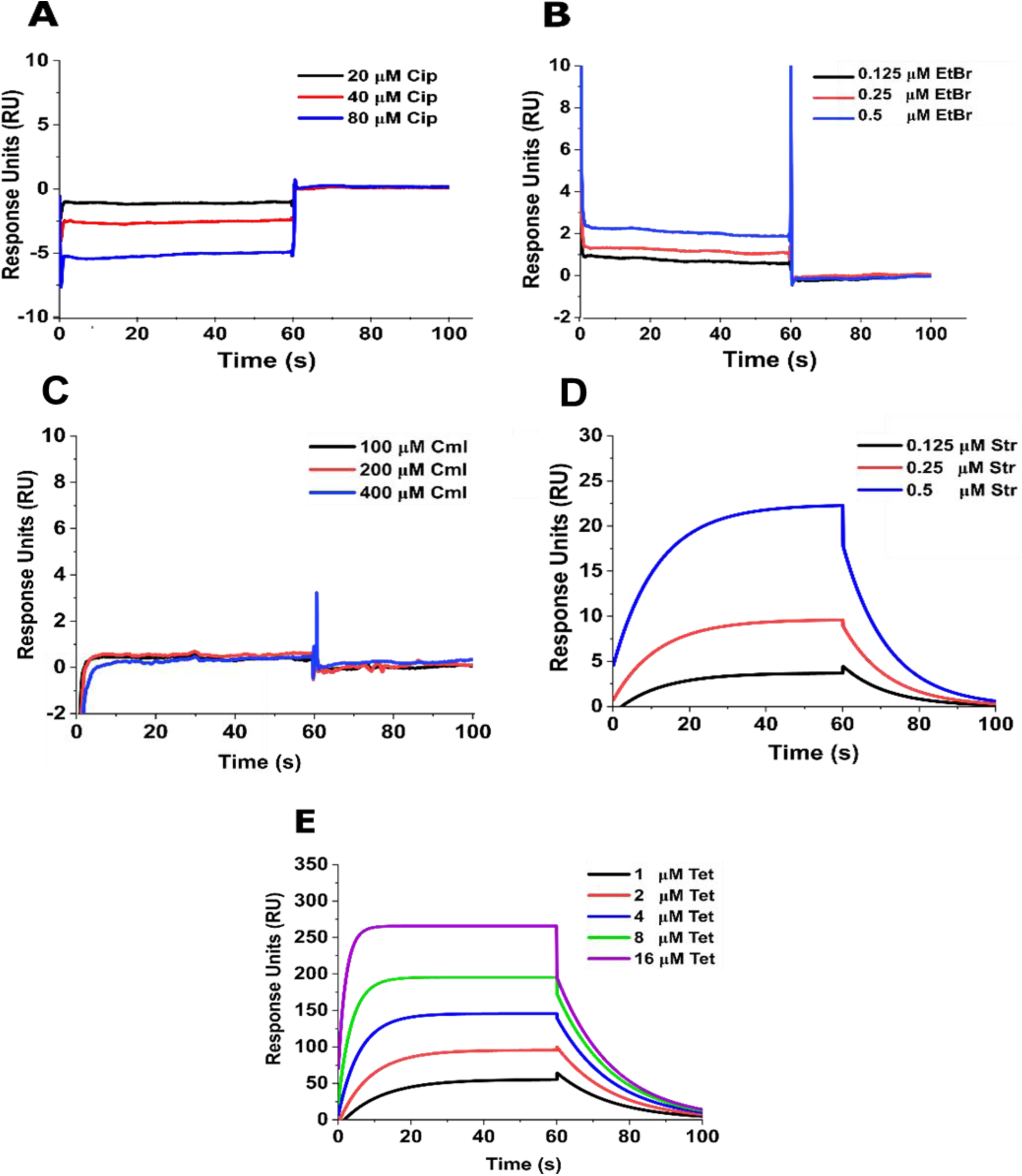
Binding kinetics of different drugs to M93L-SCO4122 mutant protein measured via SPR assay (A) Ciprofloxacin (B) EtBr (C) Streptomycin (D) Chloramphenicol (E) Tetracycline. M93L-SCO4122 protein was immobilized over CM5 Sensor chip through Amide linkage. All drugs (ciprofloxacin, EtBr, streptomycin, chloramphenicol and tetracycline) were passed over the immobilized M93L-SCO4122 protein at a concentration dependent manner. Equilibrium Dissociation constant (K_D_) of the binding of the drugs to the mutant M93L-SCO4122 protein is calculated using the Biacore software. No binding was observed for ciprofloxacin, chloramphenicol and EtBr. Streptomycin bound to the M93L-SCO4122 protein with a K_D_ of 53.9 µM. Binding of all drugs with M93L-SCO4122 was studied using an association time of 60 seconds followed by a dissociation time of 120 seconds.

### Interaction of different drugs with SCO4122 leads to enhanced affinity of SCO4122 to the *sco4121* promoter

The in-vitro binding studies using SPR, show that SCO4122 binds to *sco4121* promoter with moderate affinity. Further, SCO4122 also displayed weak affinity of binding to the various drugs alone, which are substrates of SCO4121 efflux pump (Figure 1A and Figure 2). We next understand the transcriptional activation of SCO4121 by SCO4122 in response to the drugs. For this, we again utilized SPR with the biotinylated DNA fragment corresponding to the intergenic region between *sco4121* and *sco4122*, consisting of the putative *sco4121* promoter. This biotinylated fragment was immobilized on the SA chip through streptavidin linkage. The purified SCO4122 protein was passed over the immobilized DNA and dissociation constant between the two was measured in absence and in presence of different drugs. Note that the three binding motifs of SCO4122, discussed earlier (Figure 1B), were intact in this intergenic region. It was noted that in absence of any drug, SCO4122 bound to the DNA with a K_D_ of 226 nM (Figure 6A). We also observed that binding of SCO4122 to the immobilized DNA was saturated beyond 2 µM. The equilibrium dissociation constant was further measured in the absence and presence of three substrates of SCO4121, which included ciprofloxacin, chloramphenicol and streptomycin. EtBr was not used in this assay because of its DNA binding property that might result in false positives. We selected 2 µM of the purified SCO4122 protein with varying amount of the drugs and monitored the equilibrium dissociation constant. We found that in presence of the drugs, SCO4122 protein bound to the *sco4121* promoter DNA with an enhanced affinity, where K_D_ was calculated to be 3.1 nM, 81.8 nM and 9.1 nM for ciprofloxacin, chloramphenicol and streptomycin respectively. These K_D_ values corresponded to nearly 10,000, 2000 and 25000 lower than that observed when the drugs alone bound to WT SCO4122 protein (Figure 6B, 6C and 6D). The K_D_ values of the interaction studies in absence and presence of drugs are provided in Table 2. Various studies have shown that interaction of drugs with the regulator affect its DNA binding activity. For example, in *B. subtilis*, drugs such as TPP and rhodamine interact with the MepR regulator BmrR which further enhances the transcription of the efflux gene *bmr* (30). Similarly, drugs such as kanamycin and ampicillin interact with TcaR, thus modulating its DNA binding activity in the gram-negative bacteria *Staphylococcus aureus* (31). The results generated from our study clearly indicate that the substrates of SCO4121 play an essential role in modulating the expression of SCO4121 by directly interacting and enhancing the affinity of the SCO4122 protein to the *sco4121* promoter. Note that in presence of tetracycline, which is not recognized as a substrate of SCO4121, there was no change in the affinity of *sco4121* promoter to SCO4122 protein (Figure 6E). We also found that under both in-vitro and in-vivo environment, mutant M93L SCO4122 protein was unable to bind to the *sco4121* promoter. It should also be noted that no change in the binding affinity of the M93LSCO4122 to the *sco4121* promoter was observed in the presence of the drugs: ciprofloxacin, streptomycin and chloramphenicol (data not shown).

**Figure 6:**
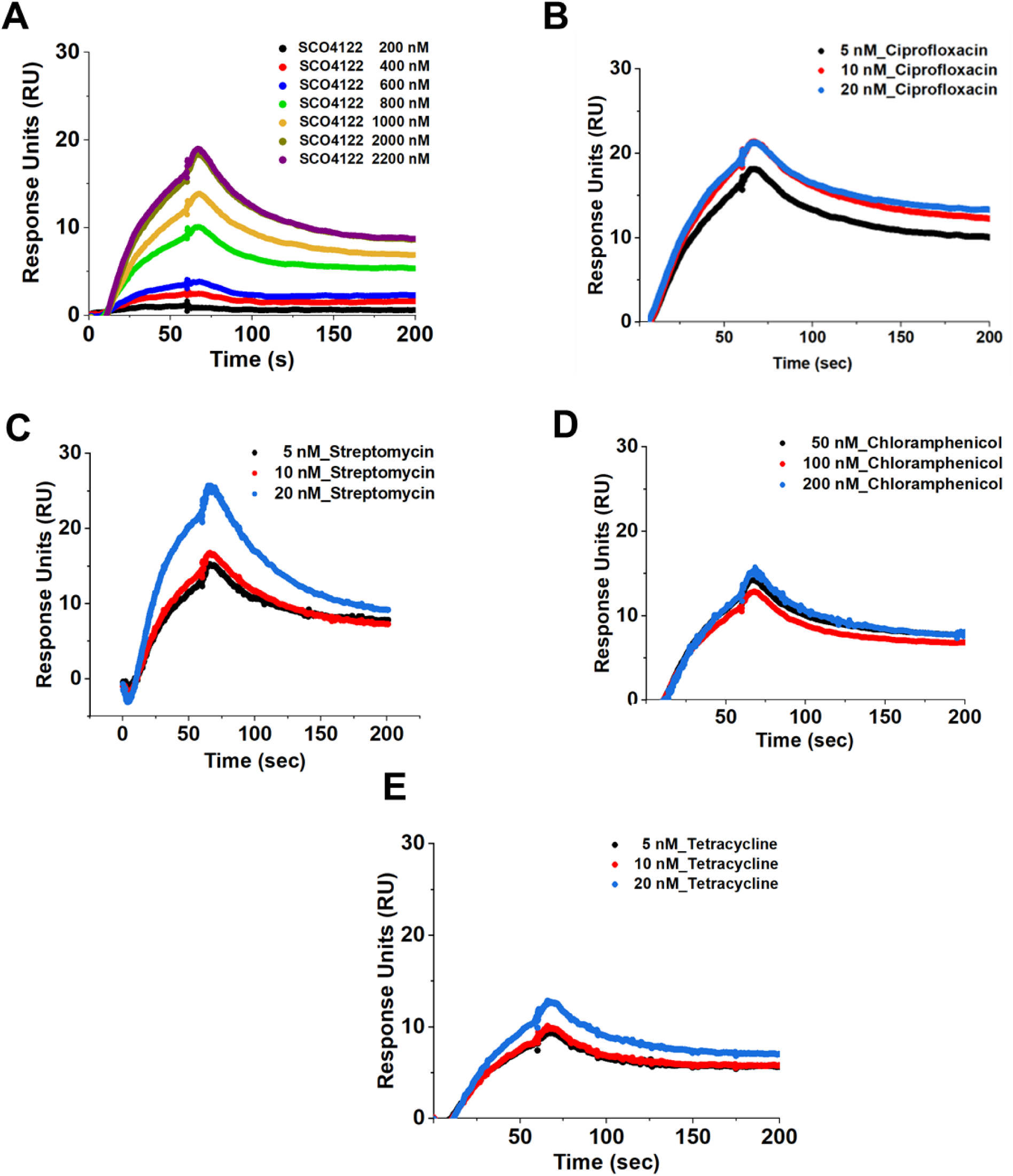
Effect of SCO4121 substrates on DNA binding ability of SCO4122 to the *sco4121* promoter. (A) Binding of purified SCO4122 protein to the biotinylated DNA of intergenic region between *sco4121* and *sco4122* containing the three motifs. The concentration of protein was varied from 200 nM to 2.2 μM. Binding of purified SCO4122 protein to the biotinylated *sco4121* intergenic region in presence of (B) 5 nM, 10 nM and 20 nM of ciprofloxacin (C) 50 nM, 100 nM and 200 nM of streptomycin (D) 50 nM, 100 nM and 200 nM of chloramphenicol. (D) 5 nM, 10 nM and 20 nM of tetracycline. Based on binding of SCO4122 with DNA, a concentration of 2 μM of protein was selected for subsequent binding experiments in presence of drug. Equilibrium Dissociation constant (K_D_) of SCO4122 with the biotinylated DNA consisting the *sco4121* promoter was increased in presence of ciprofloxacin, chloramphenicol and streptomycin. No change in K_D_ was seen in case of tetracycline. Binding was studied using SPR assay with a flow rate of 30 µl/min with an association and dissociation time of 60 seconds and 300 seconds respectively.

**Table 2:**
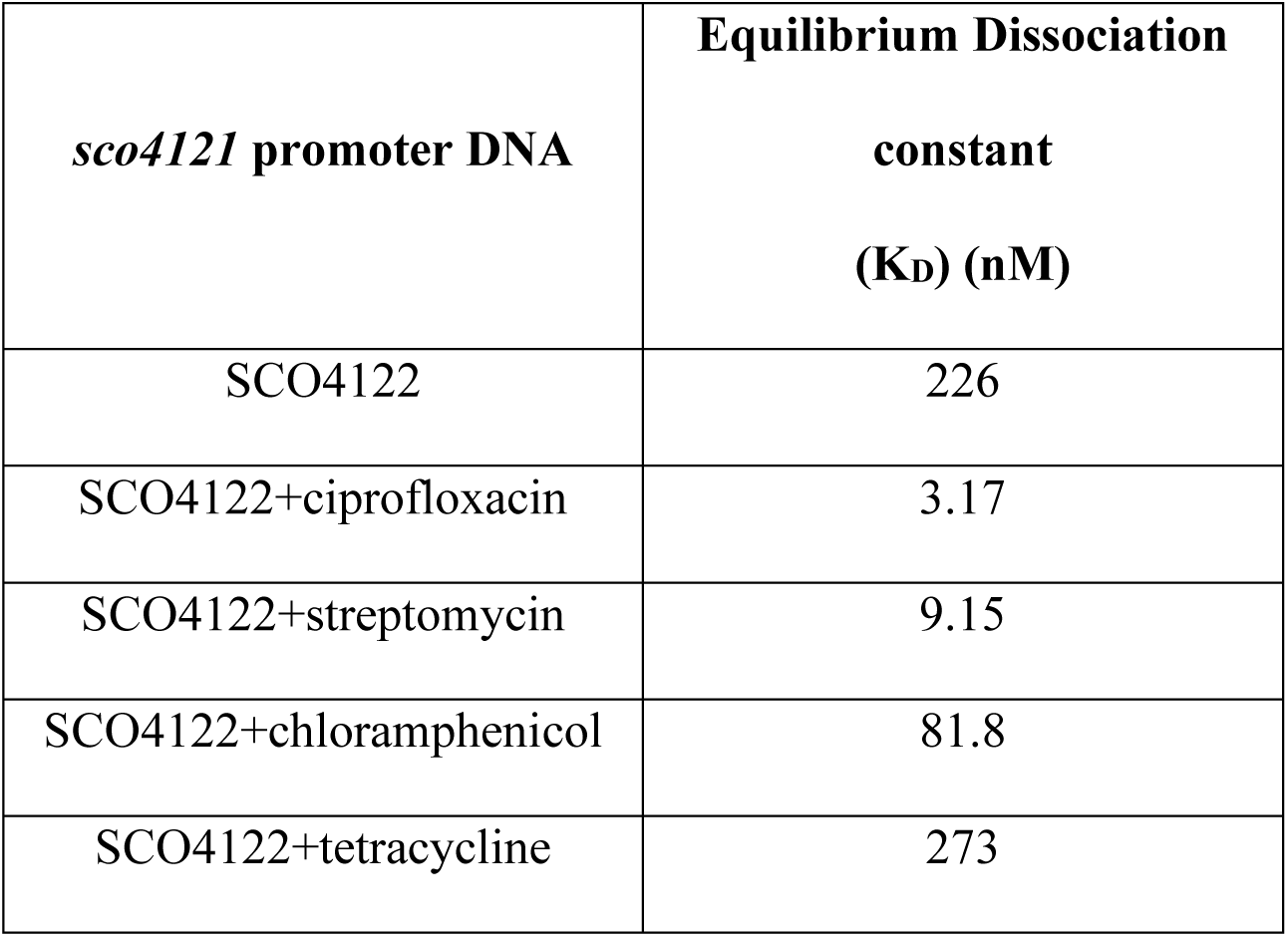
Equilibrium Dissociation constant (K_D_) of interaction between SCO4122 with the *sco4121* promoter DNA in presence of different drugs.

## Discussion

Expression of efflux pumps is finely tuned by transcriptional activators or repressors (2, 7). Both two-component and one-component transcriptional regulators have been found to be involved in regulation of efflux pumps that confer multidrug resistance phenotype in different bacteria. The MarR group of transcriptional regulators is one of the widely studied one-component regulators that are ubiquitously distributed in different bacteria and regulate a diverse array of efflux pumps (33). Previous studies in our lab, using in-vivo approaches, led us to the identification of a MarR regulator, SCO4122, in *Streptomyces coelicolor* that was associated with transcriptional activation of the efflux pump, SCO4121 in response to multiple drugs that were substrates of SCO4121 (26). In this study, using SPR, we identified that SCO4122 directly interacts with the substrates of SCO4121 with varying affinities, thereby leading to activation of the latter. Streptomycin bound to SCO4122 with the highest affinity, followed by EtBr, ciprofloxacin and chloramphenicol respectively. Our study further revealed that in response to the drugs, SCO4122 bound to three individual motifs in the *sco4121* control region. This binding was likely to be cooperative in nature as absence of any one of these motifs did not induce the expression of SCO4121, even in the presence of drugs. Transcriptional regulators have previously been shown to undergo cooperative binding to ensure effective transcription of the target gene. This regulation can be either via protein-protein interaction or through modulation of DNA topology (34). For example, the MarR regulator Rdh2R has been found to act cooperatively as a tetramer for efficient repression of *rdh* genes in *Dehalococcoides mccartyi* (35). In *Bacillus subtillis*, cooperative binding of the MarR regulator OhrR to two inverted half-sites increases the affinity of interaction in the *ohrA* control region, thus leading to its effective repression (36). Also, in presence of the inducer cadmium, the MepR regulator CadR cooperatively binds to two sites in the promoter of the efflux pump CadA, which further operates to export cadmium from the internal to the external environment (37). On similar lines, we hypothesize that in response of different substrates of SCO4121, cooperative binding of SCO4122 to the three motifs in the *sco4121* control region results in increased expression of SCO4121.

Our studies further revealed that upon interaction with different drugs, SCO4122 binds with higher affinity to the *sco4121* promoter, thus leading to enhanced expression of SCO4121. Previously, studies have shown that the MerR regulator BmrR in *Bacillus subtilis* interacts with the substrates of Bmr, thereby leading to enhanced transcription of the efflux pump Bmr (38). This interaction has been found to depend upon the conformational changes that occur in drug bound BmrR which leads to modulation of spacer sequences between −35 and −10 regions of the *bmr* promoter. This DNA distortion leads to effective recruitment of RNA polymerase and enhanced expression of Bmr (39). We hypothesize that in addition to cooperative binding, a plausible conformational change in SCO4122 in response to the different drugs results in effective binding to the *sco4121* promoter, thereby promoting its enhanced expression.

A third part of our study highlighted on the importance of a conserved methionine residue M93 in SCO4122 to play a critical role in activation of SCO4121 in response to the different substrates of SCO4121. Our previous studies have highlighted that overexpression of SCO4122 resulted in increased expression of SCO4121when induced with the substrates of SCO4121, and subsequently led to elevated resistance towards these compounds (26). In this study, under in-vivo conditions, we found that when methionine M93 in SCO4122 was converted to Leucine, *sco4121* levels remain similar to basal levels and were not upregulated in response to ciprofloxacin, streptomycin and EtBr. This also correlated with the resistance profile where M93L-4122 strains displayed similar resistance towards different drugs as that of WT86 cells. However, it was interesting to note that in presence of chloramphenicol, *sco4121* levels in M93L-4122 were upregulated similar to what was noticed for cΔ4122 cells. Along with upregulated *sco4121* levels, the M93L-4122 cells showed elevated resistance towards chloramphenicol which was also similar to that of cΔ4122 cells. However, under in-vitro conditions M93L SCO4122 protein showed negligible binding to ciprofloxacin, EtBr and chloramphenicol and bound to streptomycin with a very low affinity which was physiologically irrelevant. Previous studies have shown methionine to be essential in structural integrity of several proteins by creating bonds with the aromatic amino acids (40, 41). This suggests that methionine M93 in SCO4122 is essential in possible interaction of the drugs ciprofloxacin, EtBr and streptomycin with the protein leading to enhanced structural stability of SCO4122, thereby allowing effective activation of SCO4121. We found that the WT SCO4122 protein displayed a 2-state binding kinetics upon binding to chloramphenicol, thereby suggesting the protein to undergo a probable conformational change in response to chloramphenicol. However, the increased expression of *sco4121* in response to chloramphenicol in M93L-4122 strains suggests that the conformational change attained in presence of chloramphenicol may not necessarily involve the M93 amino acid, thus its inactivation has no impact on either the expression levels of *sco4121* or its susceptibility towards the drug. Previous studies have shown that certain teicoplanin derived antibiotics display anti-cooperativity among ligand binding and dimerization, where binding of ligand allows the protein to attain a closed conformation (42). On similar lines we hypothesize that chloramphenicol binding to SCO4122 may lead to a closed conformation which does not involve methionine M93.

In addition to this, methionine in certain transcriptional regulators has been found to be involved in sensing oxidative stress, thereby preventing bacterial killing when subjected to ROS generating compounds. One such example is the LysR regulator HypT in *E. coli*, where involvement of three methionine residues have been reported to sense oxidative stress in response to the strong oxidant HOCl, thus protecting the bacteria against it. Homologs of HypT in *Salmonella. typhimurium* have also been found to protect the bacteria against oxidative stress through oxidation of methionine (43). Our previous studies have highlighted that SCO4122 induces increased expression of SCO4121 in presence of diverse ROS generating drugs. This expression is further quenched to basal levels upon addition of an antioxidant ascorbate (26). Studies have shown that under in-vivo environments, oxidation of transcriptional regulators results in increased structural stability, thereby leading to efficient transcription of cognate genes. For example, in *Bacillus subtillis*, the transcriptional regulator Spx, upon oxidation aids RNA polymerase to form an active complex with the promoter DNA, thereby allowing transcriptional activation of several genes (44). We thus hypothesize that under in-vivo environments, oxidation of Methionine M93 can be an important factor for activation of SCO4122 and subsequent activation of SCO4121. These results thus suggest that methionine M93 could be plausibly associated in maintaining the functional conformation of SCO4122 when exposed to the substrates of SCO4121. This could be either via oxidation of methionine residues or through an unknown mechanism that helps in subsequent activation of SCO4121. We also hypothesize a plausible allosteric modification of SCO4122 in response to chloramphenicol to be sufficient to activate SCO4121, without aid of methionine M93.

Our data shows that chloramphenicol binds with the weakest affinity to WT SCO4122 protein. Additionally, upon interaction with chloramphenicol, the dissociation constant (K_D_) of WT SCO4122 to the *sco4121* promoter changes from 2000 nM to 80 nM. Despite the large change in binding affinity, the K_D_ was significantly higher as compared to that of SCO4122 binding to the *sco4121* promoter in presence of ciprofloxacin or streptomycin. Earlier, we have seen that under in-vivo environments, SCO4122 overexpression leads to upregulation of *sco4121* to similar levels in presence of ciprofloxacin, chloramphenicol and streptomycin (26). This suggests that under in-vivo environments, activation of SCO4121 by SCO4122 in presence of chloramphenicol may be coupled to an unknown mechanism apart from its direct interaction with SCO4122, which may result in further increase in the binding affinity of SCO4122 to the *sco4121* promoter. We predict ROS generated in response to chloramphenicol may be an essential factor for activation of SCO4122 that may potentially involve a conformational change through oxidation of other conserved methionine residues such as M44, M100 and M134.

Thus, taken together our study highlighted the intricate regulatory mechanisms of SCO4122 involved in regulation of the efflux gene, SCO4121. MarR regulators have previously been shown to be induced in response to a diverse array of compounds. Salicylate and copper ions have been found to be potent inducers for the MarR regulator in *E. coli* (14, 20, 45). Also, many other MarR regulators have been found to be active under redox sensing environments upon exposure to antibiotics or cellular metabolites (22, 46, 47). However, the direct interaction of different drugs with MarR regulators and their possible effects on gene expression has been poorly studied. This work demonstrates that the MarR regulator SCO4122 interacts with the substrates of the efflux pump SCO4121 in a direct manner and promotes enhanced transcription of efflux pump gene via binding to the *sco4121* promoter with high affinity. The data also suggests that the regulation of SCO4121 by SCO4122 follows a cooperative binding to three distinct motifs in the *sco4121* control region when exposed to multiple drugs. Further work is required to delineate the mechanism of cooperative binding. Furthermore, our studies also identified that under in-vivo environments, the conserved Methionine M93 plays a crucial role in functional conformation of SCO4122 which further activates SCO4121 when exposed to multiple substrates.

Several MarR regulators across bacteria have been studied that regulate gene expression via directly binding to promoter of different genes. Our results showed SCO4122 binds to *sco4121* promoter with a high dissociation constant of 0.4 µM, suggesting weak affinity as compared to that observed for many other MarR regulators binding to the promoter of their cognate genes. For example, MepR in *Staphylococcus aureus* has been found to bind to *mepR* DNA with an affinity of 36 nM (48). Also, MosR and HypS in *Mycobacterium tuberculosis* abound to its cognate DNA with an affinity of 15 nM and 70 nM respectively (47, 49). However, in presence of drugs, the binding affinity to DNA increases many folds. For example, SCO4122 bound to the *sco4121* promoter strongly with a K_D_ of 3 nM when incubated with ciprofloxacin, thus displaying an affinity remarkably higher than other MarR regulators. Comparable affinity of binding has been previously reported in case of another MarR regulator, SCO3205 from *S. coelicolor*, which binds to dual operator sites of *sco3204* with an affinity of 1.3 nM and 2.4 nM respectively and represses its transcription (50). However, our study highlights SCO4122 to be a strong activator of SCO4121 that is tightly regulated by multiple drugs leading to conditional activation of the latter. Our study thus highlights the quantitative estimates of the change in binding affinity when a bacterium senses dissimilar drugs in the environment. This model can further be utilized to understand how an efflux pump can be positively regulated in an environment where multiple stressors are available. Further biochemical and structural studies will also help us to identify whether monomeric or dimeric forms of SCO4122 involve in cooperative binding, thus aiding SCO4121 activation. Additionally, structural studies governing the interaction of SCO4122 with the drugs and the SCO4121 promoter region will provide further insight into regulation of efflux pumps by MarR regulators that can be relevant in multidrug resistant clinical isolates.

## Materials and Methods

### Bacterial growth, strains and media conditions

*Streptomyces coelicolor* WT86 and recombinants were grown in R5 media (1% wt/vol of MgCl_2_, 10% sucrose, 1% glucose, 0.003% K_2_SO_4_, 0.002% Casamino Acids, 0.5% yeast extract, 0.5% TES buffer, trace elements) for all growth conditions and assays. Mannitol Soya Agar (Mannitol-2%, soya-2%, agar-2%) was used for preparation of spores of *S. coelicolor*. *Mycobacterium smegmatis* WT and all recombinants were grown in Luria Bertani (LB) media supplemented with 0.05% Tween 80 for all growth conditions. *E. coli* WT and recombinant cells were grown in LB media for all growth conditions. *S. coelicolor* was grown at 30°C for all assays, except for selection of knockout mutant, where temperature was shifted to 39°C. *M. smegmatis* and *E. coli* were grown at 37°C for all performed assays.

### Construction of SCO4122 mutants in S. coelicolor, M. smegmatis and E. coli

Overexpression, knockout and complementation of *sco4122* in *S. coelicolor* were carried out as mentioned previously (26). To generate a directional clone in *S. coelicolor*, *sco4122* was amplified from *S. coelicolor* genomic DNA using primers listed in Table S2. The amplified product was further digested with the restriction enzymes *Bam*HI and *Hind*III and ligated into the plasmid pIJ86 thus generating O4122. pIJ86 has bears the modified promoter p*Erm*E*.(derived from *Streptomyces erythreus*) and is found to be a replicative vector. The constructs obtained were further transformed into *S. coelicolor* via conjugation using the *E. coli* strain ET12567 (pUZ8002). R5 plates having apramycin (25 µg/ml) and nalidixic acid (50 µg/ml) were used for the selection of the transconjugants. For cloning into *M. smegmatis*, *sco4122* was amplified from genomic DNA of *S. coelicolor* using primers Listed in Table S2. The PCR product was further digested using the enzymes *Bam*HI and *Hin*dIII and ligated into the integrative plasmid pSTki, previously digested with the same enzymes to yield MS4122. pSTki is Mycobacterium specific integrative plasmid with the *Pmyc* promoter. For overexpression of *sco4122* in *E. coli*, *sco4122* was amplified from *S. coelicolor* genomic DNA using primers listed in Table S2. The PCR product was further digested with the enzymes *Bam*HI and *Hin*dIII and ligated in to the replicative plasmid pET28a under the T7 promoter to yield the construct OE4122. Kanamycin at 25 µg/ml and 50 µg/ml was used for selection of MS4122 and OE4122 clones.

Deletion of *sco4122* from *S. coelicolor* genomic DNA was performed as described previously (26). The upstream and downstream sequence with respect to SCO4122, corresponding to 201 bp and 318 bp respectively were amplified from *S. coelicolor* genomic DNA using primers listed in Table S2. The upstream sequence was digested with the enzymes *Eco*RI and *Bam*HI, whereas the downstream sequence was digested with the enzymes *Xba*I and *Hin*dIII. The digested fragments were further ligated to pKC1139 previously digested with the same enzymes. The constructs were cloned into the *E. coli* strain ET12567 (pUZ8002) and transformed into *S. coelicolor* cells via conjugation to form the knockouts Δ4122. Following incubation at 30°C for 48 hours, the MSA agar plate was flooded and 500 µg/ml of nalidixic acid (Himedia) and 1 mg/ml of apramycin (Merck). After incubation, the plates were shifted to 39°C. pKC1139 is an *E. coli-Streptomyces* shuttle plasmid having a temperature sensitive replicon which halts growth of *S. coelicolor* when temperature reaches beyond 34 °C (54). The *S. coelicolor* single over mutants were allowed to undergo further three rounds of sporulation in absence of any antibiotic to generate the double cross over mutants, which were sensitive to apramycin. All *S. coelicolor* strains used in this study has been listed in Table S3.

### Preparation of M93L SCO4122 variants in *S. coelicolor, M. smegmatis* and *E. coli*

To perform site directed mutagenesis of SCO4122 Methionine 93 to Leucine, the clones O4122, MS4122 and OE4122 bearing the WT copies of *sco4122* were utilized as templates. To generate the mutated *sco4122* constructs, PCR amplification was carried using Phusion Polymerase (Thermo Scientific) for 18 cycles using the mutation primers Listed in Table S2. The generated PCR product was further digested with the restriction enzyme *Dpn*I to eliminate the methylated parental strands. The mutations in these constructs were further confirmed via sequencing. These constructs were further transformed into Δ4122 strain of *S. coelicolor*, and WT strains of *M. smegmatis* and *E. coli* to yield the strains M93L-4122, M93L-MS4122 and M93L-OE4122 respectively. The information on these strains is listed on Supplementary Table S3.

### Preparation of luciferase constructs in *M. smegmatis*

Motif 1, Motif 2 and Motif 3 were individually amplified from *S.coelicolor* genomic DNA and placed under the *Pmyc* promoter in a modified pMpV27 called pMpV27M, where the downstream gfp reporter was replaced by luciferase gene. These strains were individually incorporated into electrocompetent *M. smegmatis* MS4122 cells, where SCO4122 was overexpressed in the integrative plasmid pSTKi, thus yielding strains MS-1, MS-2 and MS-3 respectively. Motif 1 and Motif 2 were amplified together from *S. coelicolor* genomic DNA using primers listed in Table S2. The amplified product was further digested with the enzymes *Bam*HI and *Eco*RV and ligated with pMpv27M previously digested with the same enzymes. These constructs were further introduced to MS4122 cells, thus yielding the strain MS-12. Similarly, Motif 2 and Motif 3 were amplified together from *S. coelicolor* genomic DNA using primers listed in Table S2, digested with enzymes *Bam*HI and *Eco*RV and ligated with pMpv27M previously digested with the same enzymes, followed by incorporation into MS4122 cells to yield the construct MS-23. Finally, motifs 1, 2 and 3 were together amplified using primers listed in Table S2, digested with enzymes *Bam*HI and *Eco*RV and incorporated into pMpV27M plasmid, previously digested with the same enzymes. These constructs were incorporated into *M. smegmatis* to yield the strain MS-123. All these strains are listed in Table S3.

### Measuring *sco4121* promoter activity

*M. smegmatis* cells harbouring the empty plasmids pSTKi along with the constructs MS1, MS2, MS3, MS-12, MS-23 and MS-123 were grown till mid-logarithmic phase and induced with ciprofloxacin, chloramphenicol, streptomycin and EtBr for 1 hour. Activity of the *sco4121* promoter was captured by measuring the expression of the luciferase gene (as described above) post induction with the different drugs. The data was normalised using 16S rRNA as the housekeeping gene followed by normalisation with WT27 cells (cells harbouring the empty pMpV27M plasmids).

### Gene expression studies

Extraction of RNA from *S. coelicolor* and *M. smegmatis* was carried out using the Trizol method as described earlier (55). After desired growth of the cells, 12% of 5% Phenol-ethanol mixture was added and stored at −80°C before further processing. The cells were freeze thawed and 1 ml of ice cold Trizol was added to the cells. RNA was further extracted using the established protocol. cDNA synthesis was further carried out using RevertAid H minus reverse transcriptase (RT) enzyme (Thermo Fisher Scientific). The reaction mixture included RNA template of 4 µg, 0.2 mM dNTPs, 0.2 µg random primers, 1X RT buffer and 1 U of RT enzyme, followed by addition of water to 20 µl. The cycle steps for cDNA synthesis were: 25°C for 10 min, 50°C for 1 h, and final extension at 72°C for 10 min. After successful synthesis, cDNA was diluted to a final concentration of 100 ng/µl. Expression studies were further carried out using 100 ng of cDNA template and 0.5 µM (each) of specific primers in a Bio Rad CFX96 real time PCR with SyBr Green iTaq Mix (Biorad). Data generated was further analysed using 2^-ΔΔCT^ method. Normalization of the data was done with the housekeeping gene, followed by the untreated sample as and when required. For *S. coelicolor*, 23S was taken as the housekeeping gene, whereas for *M. smegmatis*, 16S was taken as the housekeeping gene. Primers used for qPCR studies for *23rRNA, 16S rRNA, sco4121, sco4122* and *luc*iferase genes were the same as used in previous work (26).

### Overexpression and Purification of SCO4122 native and mutant proteins

*E.coli* BL21 (DE3) cells harboring WT (OE4122) and M93L mutant *sco4122* (OEM4122) genes were grown in LB broth till OD reached ∼0.5. The cells were then induced with 1 mM IPTG (HiMedia) and incubated at 37°C for 4 hours. The induced cells were further pelleted by centrifuging at 6000rpm for 10 min at 4°C. The pellet was then resuspended in 20ml resuspension buffer (25mM HEPES pH 7.4, 100mM NaCl, 0.3% Sarcosyl (HiMedia), 1mM PMSF) and sonicated at 40 amplitude for 20 minutes with 1sec ON and 1 sec OFF cycle. The cell lysate was centrifuged at 12,000 rpm for 40 min at 4°C and supernatant was added to 2ml of equilibrated Ni-NTA beads (Gen Script, USA Cat no-L00223-10). Equilibration buffer (25mM HEPES pH 7.4, 100mM NaCl) was passed over Ni-NTA beads for 2h at 4°C to equilibrate them. The cell supernatant was further loaded onto the equilibrated Ni-NTA beads, and flow-through was collected. The protein bound beads were then washed with 15 bead volumes of wash buffer-1 (25mM HEPES pH 7.4, 100mM NaCl) and 10 bead volumes of wash buffer-2 (25mM HEPES pH 7.4, 100mM NaCl, 10mM Imidazole). Elution of the protein was carried out using elution buffer (25mM HEPES pH7.4, 100mM NaCl, 100mM Imidazole). The eluted proteins were further concentrated using Amicon^®^ Ultra-4 centrifugal filter (Millipore) and contaminants were removed using Gel-filtration chromatography. The WT SCO4122 protein was purified via passing through the column (Superdex^TM^ 75, Biorad) at a flow rate of 0.125ml/min and pressure of 76-77psi, whereas the mutant M93L SCO4122 was purified through the column Sephadex 200 (AKTA) by passing at a flow rate of 0.75 ml/min. The subsequent peaks were collected and checked by running on 10% SDS PAGE. Cells harbouring the empty plasmid pET28a were used as negative control in this study.

### Western Blotting

For visualization of samples through Western blot, the WT SCO4122 and mutant M93L-SCO4122 proteins were transferred on to polyvinylidene difluoride (PVDF) membrane (Millipore) using transfer buffer (6g Tris base, 2.88g Glycine, 30ml methanol and final volume to 200ml with autoclaved DDW) in a semidry transfer unit (Genei) at 250mA, 40V for 45min. The blot was further blocked using blocking buffer [5% Skimmed milk powder (Hi Media) in 1X TBST] for 1 hour at 37 °C. After removal of blocking buffer, the blot was incubated with Primary antibody [Mouse anti -His tag antibody (Cat No: AE003, Abclonal)] overnight at 4°C. The primary antibody was further removed by washing the blot thrice with 1X TBST and incubated with Secondary antibody [Goat anti-mouse IgG HRP conjugated (Cat No: AS003, Abclonal)]. After removal of secondary antibody, the blot was washed thrice with 1X TBST with and developed by adding the substrate 3,3’, 5 5’ Tetramethylbenzidine (TMB, Hi Media). Figure S7 shows the expression of WT SCO4122 and M93L SCO4122 protein upon detection with Ani-histag antibodies.

### Antibiotic susceptibility assays

The antibiotic susceptibility of *S. coelicolor* WT and recombinants was done using Disc Diffusion assay. 10^8^ spores of *S. coelicolor* were plated onto R5 plates with apramycin at a concentration of 25 µg/ml. Sterile Discs (Himedia) impregnated with the desired antibiotic was placed at equidistant positions on the plates. The plates were further incubated for 72 hours at 30°C. Zone of inhibition was measured after the incubation period. The higher zone indicated increased susceptibility towards the drug. The drugs ciprofloxacin, chloramphenicol, streptomycin and EtBr were used for the assay. Ciprofloxacin and apramycin was purchased from Sigma-Aldrich whereas all other drugs were purchased from Himedia.

### Surface Plasmon Resonance Assay

#### Protein-DNA/drug interaction

The interactions of the wild type and mutant SCO4122 protein with DNA and other drugs were measured using the SPR method (Biacore X100 system, GE Science) operating at a flow rate of 30μl/min. This is a label free optical method to study the binding of different biomolecular interactions. Both WT and mutant M93L SCO4122 proteins were separately immobilized into the Carboxymethyl dextran (CM5) sensor chip using the Amine coupling. The CM5 sensor chip was activated using the mixture of 1 ethyl-3-(3-dimethylaminopropyl) carboiimide and N-hydroxysuccinimide (1:1). For SPR studies, 3 µg of pure SCO4122 protein was immobilized into the CM5 sensor chip reaching an initial RU value of 3000; whereas for M93L mutant protein, an initial RU of 1400 was detected which corresponded to immobilization of 1.5 µg of the protein. For each of the conditions tested, the DNA and drugs were passed over WT and M93L mutant SCO4122 protein at different times and at varying concentrations. The association time of interaction was fixed at 60 seconds, followed by a dissociation time of 120 seconds. K_D_ of the interactions was measured using the data from Biacore software. The parameters for the fitting were the global 1:1 binding except in the case of chloramphenicol where two-state fitting was done. 25 mM HEPES was used as the running buffer and for dilution of samples for all assays.

#### DNA-Protein-Drug interaction

The ssDNA intergenic region of 145bp between *sco4121* and *sco4122* (*sco4121* control region consisting the *sco4121* promoter) was taken and biotinylated at the 5′ end (IDT technology, Mumbai). About 1μM of biotinylated DNA was immobilized into the Streptavidin (SA) chip attached to a carboxymethyldextran. The RU value reached during immobilization was 150 RU. The mutant M93L SCO4122 protein at a concentration of 100 nM-30000 nM was passed over the biotinylated DNA. Drugs at different concentrations were incubated with 2 µM of mutant SCO4122 protein for 10 minutes and passed over the immobilized DNA. The mixture was passed with a flow rate of 30 μl/min with an association time of 60 seconds followed by a dissociation time of 300 seconds. K_D_ of each of these interactions was measured thereafter using the data using the Biacore software. 25 mM HEPES was used as the running buffer for the assay. The interaction studies were performed in duplicates and data from one of the replicates has been shown.

### Motif analysis

To determine the different motifs to which SCO4122 binds, the intergenic region consisting of the putative *sco4121* promoter was taken and a Multiple Sequence alignment was carried out with different Streptomyces species where homologs of SCO4121 and SCO4122 were conserved with greater than 90% identity. These Streptomyces species have been identified in our previous work (26). Three different motifs were predicted using the MEME tool (56). Each of these motifs showed an occurrence with a p-value of less than 0.0001. BPROM analysis tool was utilized to detect the putative promoters of *sco4121* and *sco4122* (57).

## Author contributions

AN and SM conceived of the project and designed the experiments. AN carried-out Motif determination, SPR assays related to WT SCO4122 protein, antibiotic susceptibility assays and all quantitative Real Time PCR assays. SS purified the M93L SCO4122 protein and carried out SPR assays related to it. AN, SM and SS performed data analysis. SR purified the WT SCO4122 protein. AN wrote the first draft of the paper and SM edited and reviewed it. All authors contributed to discussion and proof read the manuscript.

## Acknowledgements

The authors thank the SPR facility at IIT Bombay. They also thank Prof. Ruchi Anand and Prof. Samir Maji, IIT Bombay for providing the FPLC system for purification of proteins. AN and SS thank Department of Science and Technology (DST-Inspire) and Ministry of Education, Govt. of India (PMRF) for their respective fellowships. SR was supported through a project funded to SM from the Department of Science and Technology (SERB No. EMR/2016/007667).

## Supporting Information

Supporting information containing figures and tables is provided separately as supplementary file.

