## Supporting information for "Insight into the regulatory mechanism of the MFS transporter, SCO4121 by the MarR regulator, SCO4122"

**Running title: Regulation of efflux pump by a MarR regulator**

**Present Address:**

Ankita Nag: Department of Microbiology and Immunology, Emory University School of Medicine, Atlanta, Georgia

Table S1: DNA sequences used for SPR study

| **Sequence** | **Remarks** |
| --- | --- |
| 5’TGCCCTGGTCACGTTCACCCTCTCTTGATCTCTCGGATCGTCCGCTCCAGCCGTTAGAGCTGAAGACTTAGAAGCTAAAGTTCGAAGGTGAAGAGTACATATCGAAGGACTTCAAAGCAAAGGAGTTGCGTGCCATACTGCGGCC 3’ | Intergenic region between *sco4121* and *sco4122* with the *sco4121* control region consisting putative *sco4121* promoter |
| 5’CTCTCCTACTCCCCCAGCTACCGCTGTGCCACTGCCACTTACTTGCCGCCCGGCAAGTTGTCTACACTGGGAAAGGTAGGCGCGTAACTAGCCGGGCGTCAAGTAAGTTTTTCGGAAAGTGCGCGGGGCAGGAGGTGCGTAGGAGGATGGGCGGCACG 3’ | Non-specific (NS) DNA (Promoter region of SCO3366 efflux pump) |
| 5’TGCCCTGGTCACGTTCACCCTCTCTTGATCTCTCGGATCGTCCGCTCCAGCCGTTAGAGCTGAAGACTTAGAAGCTAAAGTTCGAAGGTGAAGAGTACATATCGAAGGACTTCAAAGCAAAGGAGTTGCGTGCCATACTGCGGCC 3’ | 5’ Biotinylated DNA with the *sco4121* control region consisting putative *sco4121* promoter used for SPR assays |

Table S2: List of primers used in this study

| **Primers (5’-3’)** | **Description** | **Reference** |
| --- | --- | --- |
| Fp LFR-ATCAGAATTCGACAGAACCCAGTCCCCGTA | Primers for deletion of SCO4122 from WT *S. coelicolor* genome | (1) |
| Rp_LFR_ AATATCTAGAGGCGATCTGCTCTTCGAGT |  |  |
| Fp_RFR_ ATATAAGCTTACTTGGGGGTCGTGCTGGTG |  |  |
| Rp_RFR-AATTTCTAGAGCGGAGGGGCGCGAGAAGT |  |  |
| Fp-AATT GGATCCGGAGTTGCGTGCCATACTGC | Primers for cloning of SCO4122 into pIJ86 in *S. coelicolor* WT and Δ4122 mutant | (1) |
| Rp-AATTAAGCTTCGCAGAGGGCTGTCAAGC |  |  |
| Fp-AATTGGATCC GGAGTTGCGTGCCATACTGC | Primers for cloning of SCO4122 into pSTKi in *M. smegmatis* | (1) |
| Rp-ATTAAAGCTT CGCAGAGGGCTGTCAAGC |  |  |
| Fp-AATT GCTAGCGGAGTTGCGTGCCATACTGC | Primers for cloning of SCO4122 into pET28a in *E.coli* BL21(DE3) | (1) |
| Rp-AATTAAGCTTCGCAGAGGGCTGTCAAGC |  |  |
| Fp-GCCGCG**T**TGACGCACCGGATCGA | Mutagenesis primers for creating the M93L SCO4122 construct under pIJ86 plasmid in *S. coelicolor ,* under pET28a in *E.coli* and under pSTKi in *M. smegmatis* | This work |
| Rp-CGGCGC**A**ACTGCGTGGCCTAGCT |  |  |
| Fp-CCTAGATATCACATATCGAAGGACTTCAAAGCA | Primers for creation of Motif 1 construct in pMpV27M | This work |
| Rp-TGCTTTGAAGTCCTTCGATATGTGATATCTAGG |  |  |
| Fp-CTTCAGCTCTAACGGCTGGAGATATCTATT | Primers for creation of Motif 2 in pMpV27M | This work |
| Rp-AATAGATATC TCCAGCCGTTAGAGCTGAAG |  |  |
| Fp-CTTCAGCTCT GAAGAGTACATATCG | Primers for creation of Motif 3 in pMpV27M | This work |
| Rp-AATAGATATGCCGCAGTATGGCACG |  |  |
| Fp-CCTAGATATCACATATCGAAGGACTTCAAAGCA | Primers for creation of Motif 1 and Motif 2 together construct in pMpV27M | This work |
| Rp-AATAGATATC TCCAGCCGTTAGAGCTGAAG |  |  |
| Fp-CTTCAGCTCTAACGGCTGGAGATATCTATT | Primers for creation of Motif 2 and Motif 3 in pMpV27M | This work |
| Rp- AATAGATATGCCGCAGTATGGCACG |  |  |
| Rp-TTAAGGATCCGGCGATCTGCTCTTCGAGTG | Primers for creation of Motif 1, Motif 2 and Motif 3 together in pMpV27M | This work |
| Fp-GCAAGAATTCATTGCCCTGGTCACGTTCAC |  |  |

Table S3: List of strains used in this study

| Strains | *Organism* | Description | Reference |
| --- | --- | --- | --- |
| M145 | *S. coelicolor* | Strain devoid of SCP1 and SCP2 plasmid | (2) |
| O4122 | *S. coelicolor* | *sco4122* cloned in pIJ86 plamid | (1) |
| Δ4122 | *S. coelicolor* | *sco4122* knocked out from *S. coelicolor* genome | (1) |
| cΔ4122 | *S. coelicolor* | Complementation of Δ4122 with WT copies of *sco4122* | (1) |
| M93L-4122 | *S. coelicolor* | Δ4122 strain consisting of M93L mutated copies of *sco4122* | This work |
| MC^2^155 | *M. smegmatis* | WT laboratory strain | (3) |
| MS4122 | *M. smegmatis* | WT MC2155 transformed with native copies of *sco4122* under pSTKi | (1) |
| MS4122:P_4121_ | *M. smegmatis* | MS4122 cells expressing *sco4121* promoter cloned under pMpV27M | (1) |
| MS-1 | *M. smegmatis* | MS4122 cells expressing Motif 1 cloned under pMpV27M | This work |
| MS-2 | *M. smegmatis* | MS4122 cells expressing Motif 2 cloned under pMpV27M | This work |
| MS-3 | *M. smegmatis* | MS4122 cells expressing Motif 3 cloned under pMpV27M | This work |
| MS-12 | *M. smegmatis* | MS4122 cells expressing Motif 1 and Motif 2 cloned under pMpV27M | This work |
| MS-23 | *M. smegmatis* | MS4122 cells expressing Motif 2 and Motif 3 cloned under pMpV27M | This work |
| MS-123 | *M. smegmatis* | MS4122 cells expressing Motif 1, Motif 2 and Motif 3 cloned under pMpV27M | This work |
| BL21 (DE3) | *E.coli* | BL21 cells with F–ompT hsdS(rB– mB–) gal dcm λ(DE3) genotype | (4) |
| OE4122 | *E.coli* | BL21 (DE3) cells expressing SCO4122 cloned under pET28a | (1) |
| OEM4122 | *E.coli* | BL21 (DE3) cells expressing M93L SCO4122 cloned under pET28a | This work |

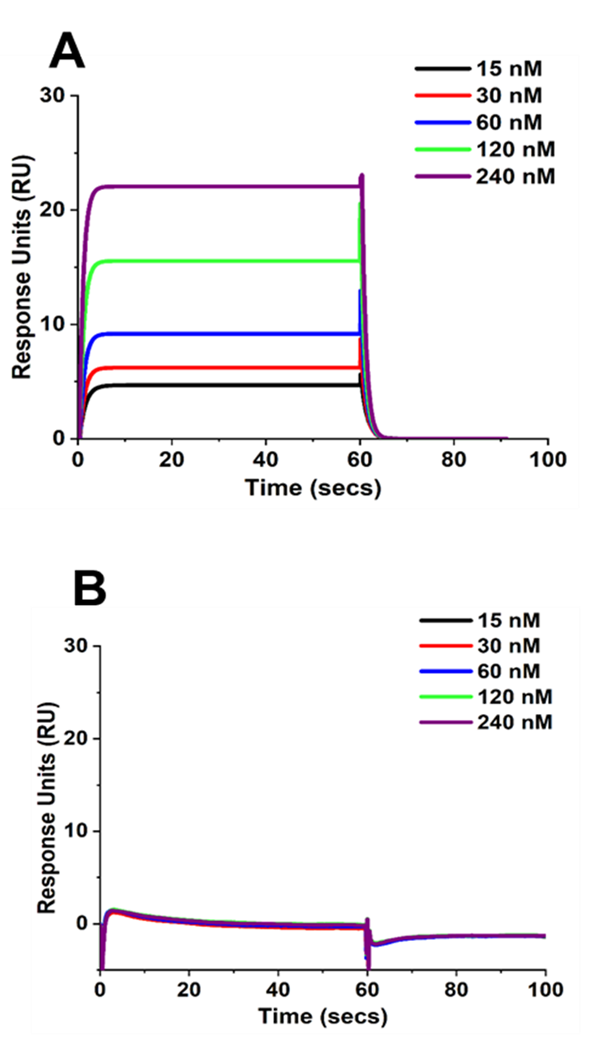

Figure S1: Binding of non-specific (NS) DNA to the purified SCO4122 protein immobilized on CM5 sensor chip using SPR assay. Binding was measured at varying concentrations of DNA with an association time of 60 seconds and a dissociation time of 120 seconds. No binding of the NS DNA to the SCO4122 protein could be seen.

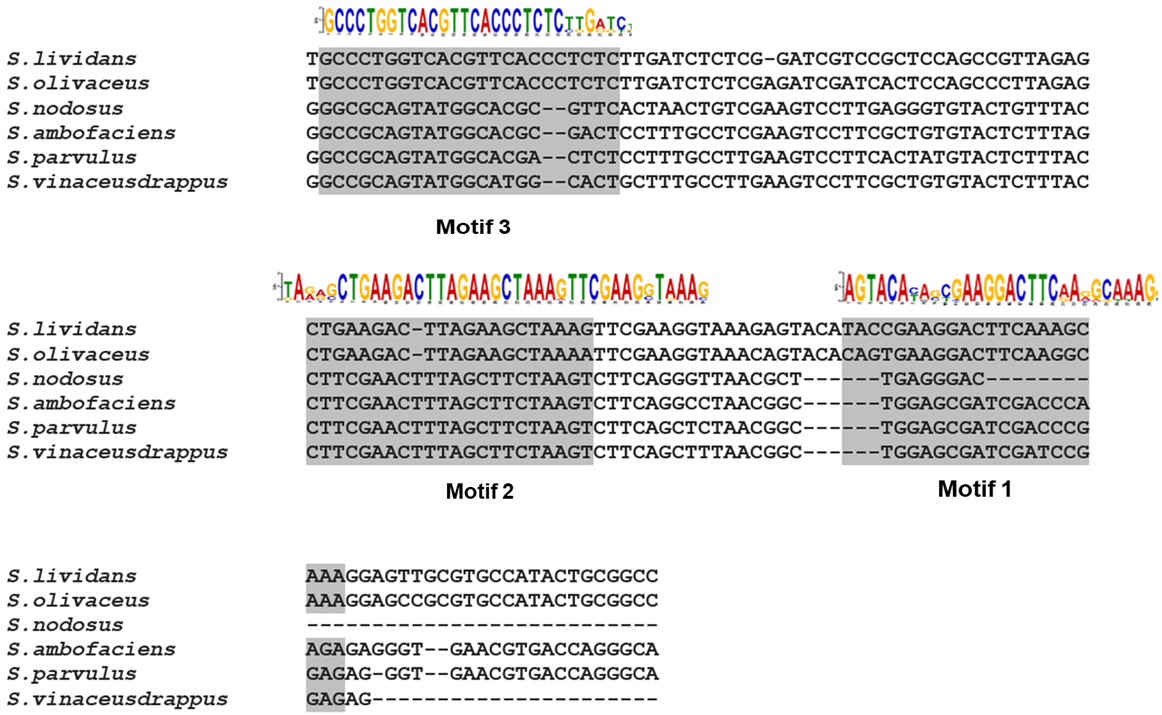

Figure S2: Distribution of Motif 1, Motif 2 and Motif 3 in different Streptomyces species where homologs of SCO4121 and SCO4122 are present with an identity of > 90%. The sequences were aligned using Multiple Sequence alignment tool CLUSTAL W and Motif search was done using the MEME tool. The conserved motifs are depicted and highlighted in the sequences

Figure S3: Schematic depicting different luciferase constructs prepared with Motif 1, Motif and Motif 3 present in the intergenic region between *sco4121* and *sco4122.* These motifs were amplified from genomic DNA of WT *S. coelicolor* and placed under the plasmid pFpV27M and transformed into MS4122 cells to generate the strains MS-1, MS-2,MS-3, MS-12, MS-13, MS-23, MS-123.
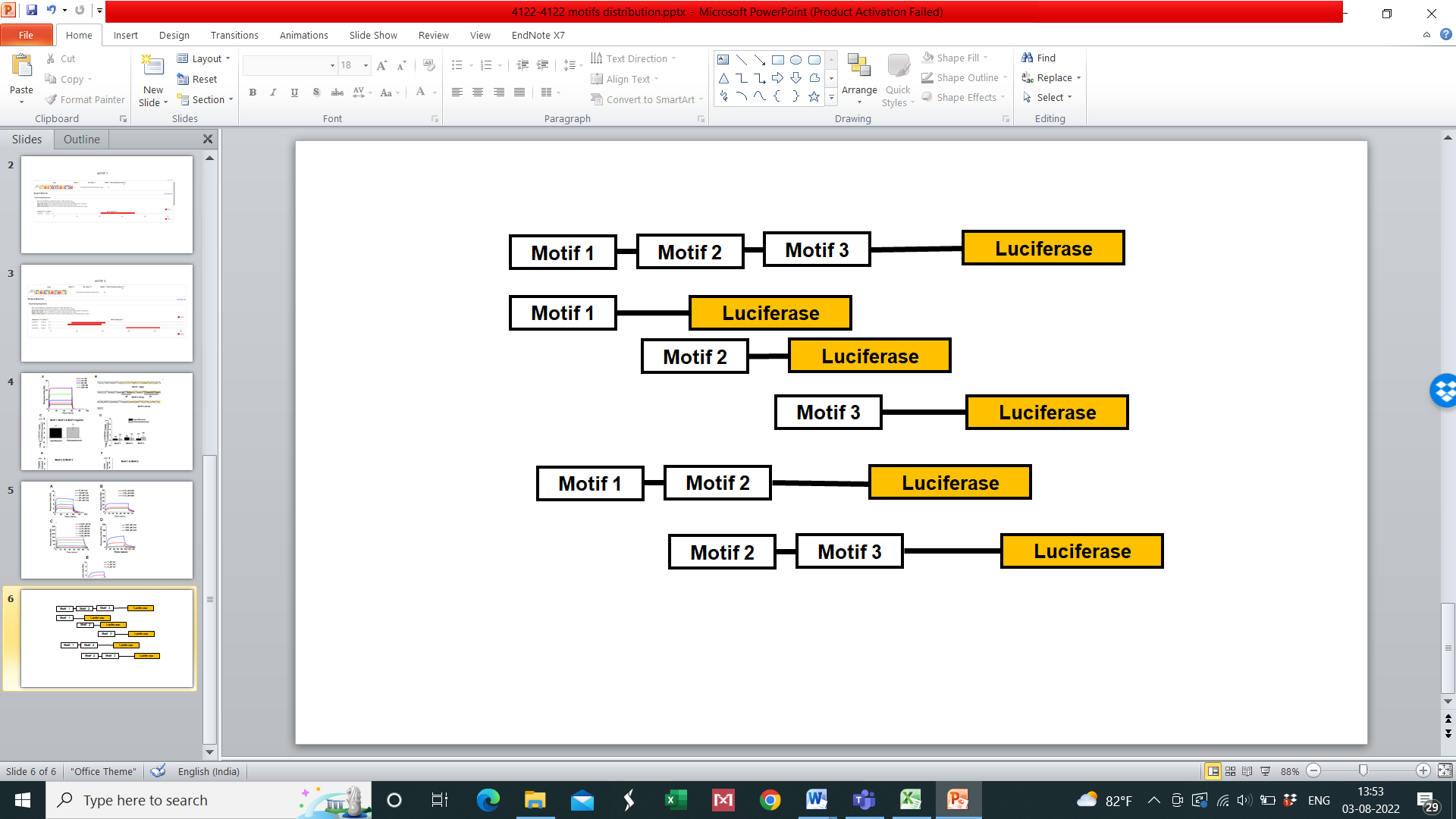

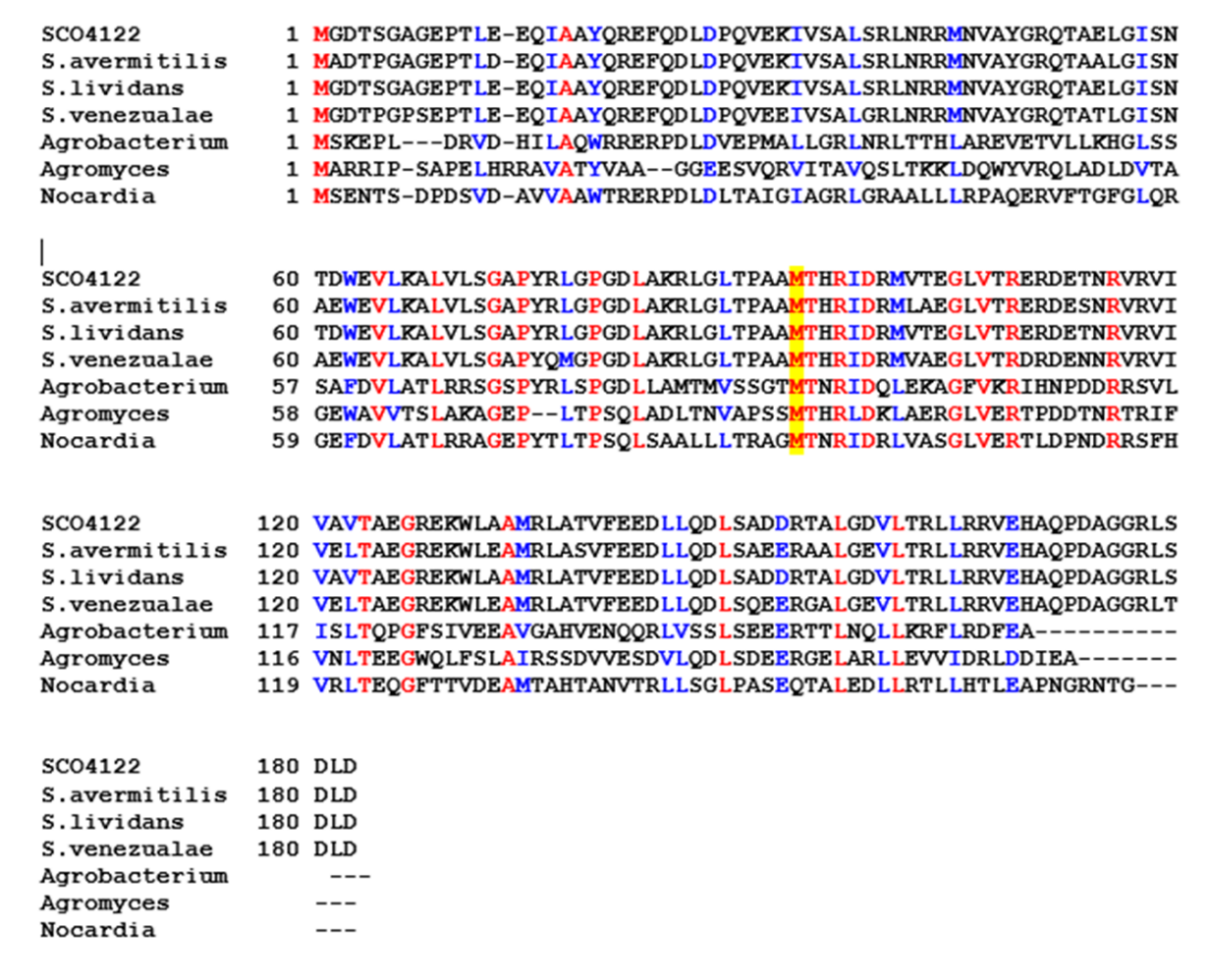

Figure S4: Multiple sequence of SCO4122 protein with protein from different streptomyces and other bacteria showing conservation of Methionine M93. This includes *S. avermitilis* (WP_010985534.1), *S. lividans* (WP_003974850.1), *S. venezualeae (*WP_055568202.1), *Agrobacterium* sp (WP_006699719.1), *Agromyces sp* (WP_189084226.1) and *Nocardia sp* (WP_068040095.1_) Multiple Sequence Alignment is performed using CLUSTAL W algorithm with T-COFFEE software.

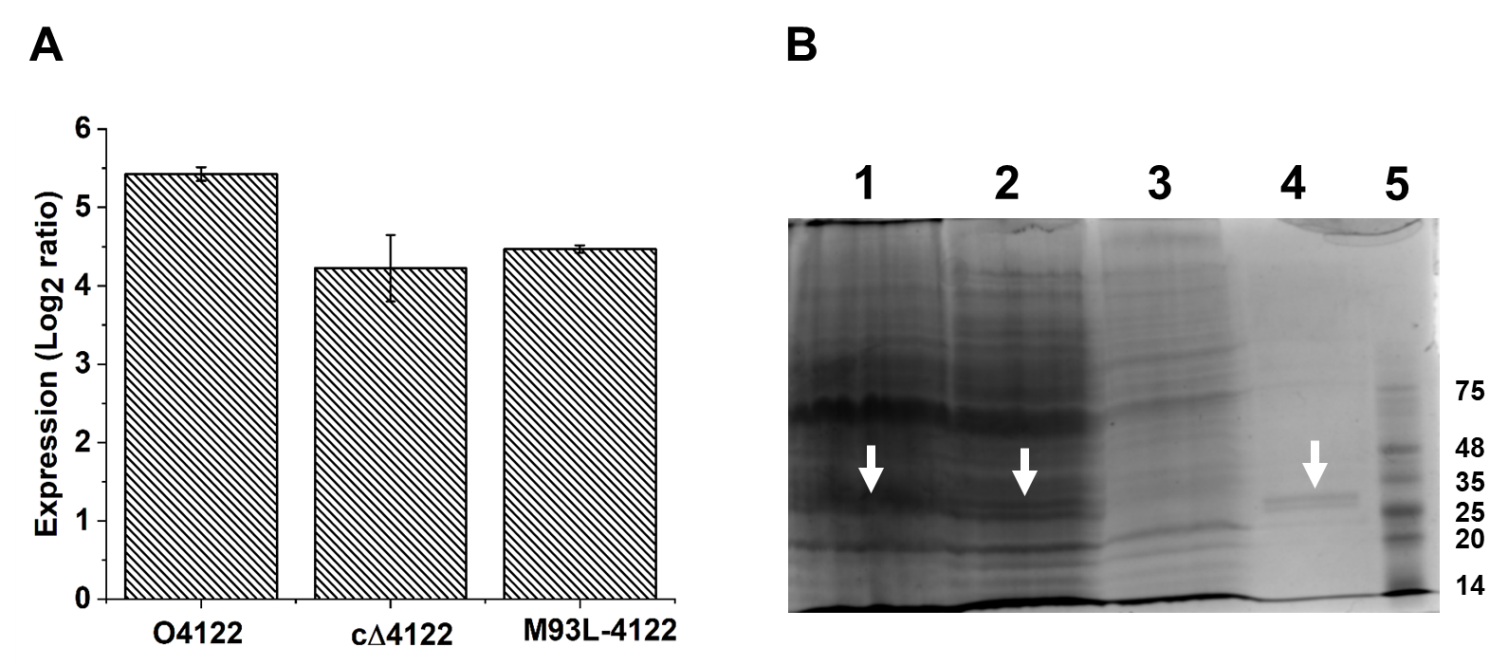

Figure S5: Expression of *sco4122* in O4122, c∆4122 and M93L-4122 strains. O4122 and cΔ4122 carried the native copies of *sco4122*, whereas M93L-4122 carried the mutated copies of *sco4122*, where the conserved Methionine M93 was converted to Leucine. Expression measured by Real Time Quantitative PCR. 23S rRNA was utilized as the house keeping gene. Expression was normalized with respect to WT cells. Expression of *sco4121* was found to be similar in all the strains

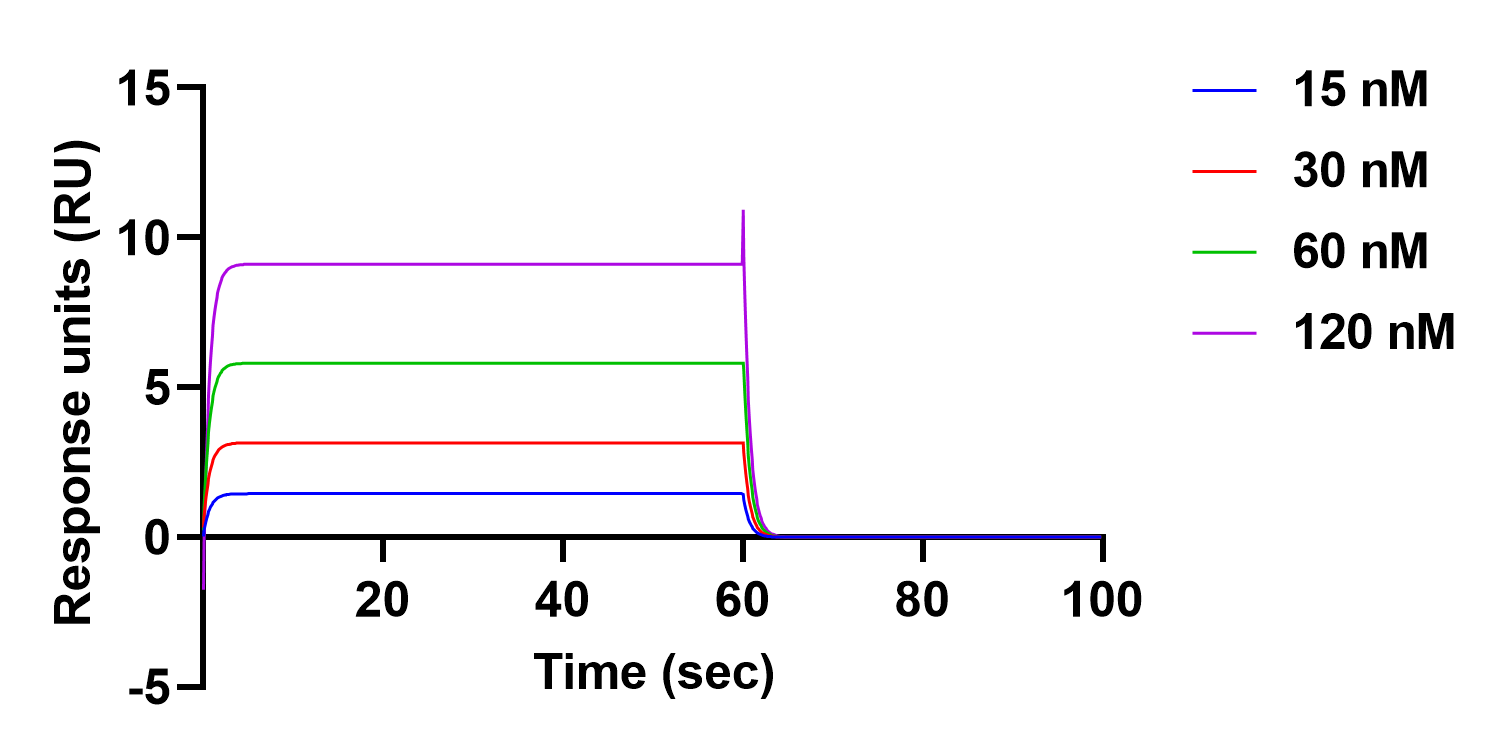

Figure S6: Binding of mutant M93L-SCO4122 protein to the *sco4121* control region consisting the *sco4121* promoter. M93L-SCO4122 was purified and immobilized on CM5 sensor chip. DNA at different concentrations was passed over the protein and binding was monitored using an association time of 60 seconds and a dissociation time of 120 seconds. M93L-SCO4122 bound to *sco4121* promoter with a K_D_ of 2.11 μM

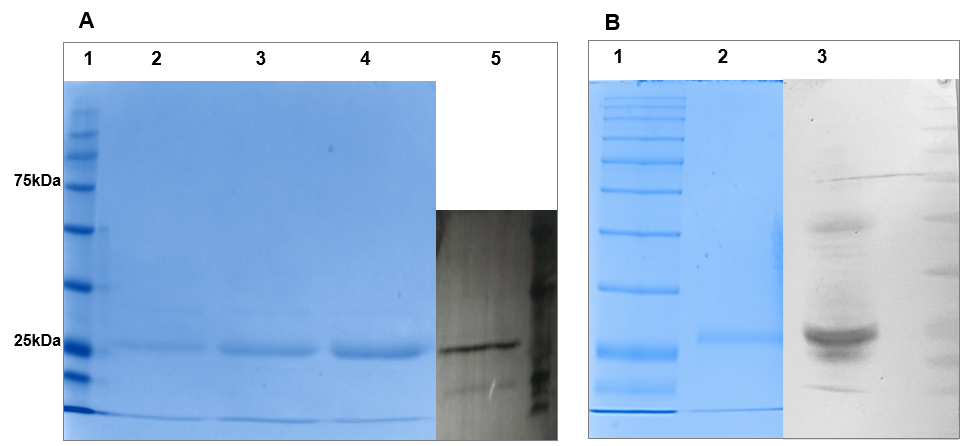

Figure S7: Expression and detection of WT SCO4122 and mutant M93L-SCO4122 protein in *E. coli* BL21(DE3) cells. (A) Expression of WT SCO4122 protein expressed in *E. coli* OE4122 cells. Lane 1: Protein marker; Lane 2, Lane 3 and Lane 4 are different Elution fractions of the purified SCO4122 detected by running on 10% SDS PAGE through, Lane 5 is the Western Blot of SCO4122 purified protein using Anti-his tag antibody. (B) Expression of M93L-SCO4122 protein in *E. coli* OEM4122 cells. Lane 1: Protein marker; Lane 2 : Elution fraction of the purified M93L-SCO4122 protein detected by running on 10% SDS PAGE, Lane 3: Western Blot of M93L-SCO4122 using Anti-his tag antibody. OE4122 and OEM4122 cells were grown in LB till mid-logarithmic phase and induced with 0.5 mM IPTG for 4 hours. SCO4122 and M93L-SCO4122 protein were extracted from these cells using the defined protocols and detected on SDS PAGE gel using Coomassie staining
